# DAZL regulates mRNA deadenylation independently of translation in germ cells

**DOI:** 10.1101/2021.02.02.429436

**Authors:** Richard W. P. Smith, Barbara Gorgoni, Zoë C. Johnston, William A. Richardson, Kelsey M. Grieve, Ross C. Anderson, Nicola K. Gray

## Abstract

Aberrant gene expression during gametogenesis is one of the factors underlying infertility, which affects roughly 15% of couples worldwide. *Deleted-in-Azoospermia-Like (DAZL),* a member of the *DAZ-gene* family, encodes an mRNA-specific regulator of translation which is essential for gametogenesis in both sexes. In this study we show that DAZL controls gene expression in oocytes by regulating the length of the mRNA poly(A) tail, a major determinant of temporal and amplitudinal gene regulation in germ cells, in which gene expression is regulated entirely post-transcriptionally. We show that DAZL does not induce polyadenylation but that binding of DAZL efficiently inhibits mRNA deadenylation induced by oocyte maturation. We reveal that this activity depends on DAZL-mediated recruitment of poly(A)-binding protein, PABP, to the mRNA. Although DAZL also activates mRNA translation via PABP recruitment, mechanistic analysis revealed that neither translation nor translational activation are required for DAZL to stabilise the poly(A) tail, suggesting two mutually independent posttranscriptional roles for the DAZL-PABP complex. We show that recruited PABP must maintain its ability to bind RNA, leading to a model in which DAZL recruits PABP and/or stabilises PABP binding to poly(A) thereby preventing access of deadenylases. These results indicate that the role of DAZL in regulating germ-cell mRNA fate is more complex than previously thought and inform on the poorly understood links between mRNA translation and deadenylation, showing that they can be mechanistically separable.

## INTRODUCTION

The level of protein production from an mRNA depends on its half-life and the efficiency with which it is translated, properties that are subject to complex regulation in all cells. It is becoming increasingly apparent that mRNA translation and stability are governed by highly interdependent processes, although in many instances the mechanistic details of their relationship remain unknown (5, 52). Central to control of both processes are numerous RNA-binding proteins (RNA-BPs) some of which can be multifunctional, participating in different regulatory complexes (45).

Removal of the 3’ poly(A) tail (deadenylation) is usually the first and rate limiting step in mRNA decay and is also associated with translational silencing or repression (10). However, recent genomewide analyses have shown that poly(A)-tail shortening is by no means always a hallmark of reduced translational efficiency, indicating that the correspondence between poly(A)-tail length and translation is more complex than previously appreciated (40, 49). Germ cells and early embryos, which are characterised by dynamic regulation of poly(A)-tail length (74), provide an excellent biological system in which to explore the links between translation and deadenylation. This has been best studied in *Xenopus laevis* oocytes, where mRNAs can undergo rapid deadenylation as they exit the nucleus, but where, in contrast to most somatic cells, this does not lead to decay but enables these mRNAs to be stored in a translationally repressed manner (57). When their products are required, translation can be activated by addition of a new poly(A) tail (cytoplasmic polyadenylation) (57), enabling the binding of poly(A)-binding proteins (PABPs) that interact with eukaryotic initiation factor (eIF)4G, a component of the 5’ cap-binding complex, bringing the 5’ and 3’ ends of the mRNA into proximity, resulting in the ‘closed-loop’ mRNA conformation that enhances cap binding and 40S-subunit recruitment (32). Cytoplasmic polyadenylation in oocytes and embryos is conserved across metazoans (53), and is also important in other cell types such as neurons (74), as well as in circadian control of gene expression (35).

*Deleted-in-Azoospermia-Like (DAZL)* is the best studied member of the *DAZ-gene* family (also comprising *DAZ* and *BOULE)* that encodes RNA-binding proteins (30, 42, 70) characterised by a conserved RNA-recognition motif (RRM) and one or more copies of the DAZ motif that mediates protein interactions (7). Members of this family are predominantly expressed in germ cells (7, 56) and are essential for germ-cell development from worms to man (17, 29, 33, 41, 60, 61, 69). DAZL and BOULE are also present in embryonic stem cells where they appear to function in exit from pluripotency during germ cell differentiation, with some studies showing they can drive germ cell development (31, 34, 82).

The molecular functions of DAZ-family members appear to be highly conserved as demonstrated by cross-species rescue experiments (7). Consistent with the importance of post-transcriptional regulation in germ cells, all tested DAZ-family proteins are mRNA-specific activators of translation initiation (14, 62). Several mRNAs that are translationally regulated by DAZL *in vivo* have been verified in both male and female mice (11, 39, 54, 55, 80) and human foetal ovary (58) with genome-wide analysis suggesting that up to 35% of mouse oocyte mRNAs may be regulated by DAZL (11). The 3’ untranslated regions (UTRs) of these mRNAs contain U-rich DAZL-binding sites that are required for DAZL to promote translation (11, 30, 54, 55, 64). We previously demonstrated that DAZL stimulates translation initiation at the 40S joining step (14, 62) and interacts directly with PABP, leading to a model in which it recruits PABP to the 3’ UTR of target mRNAs via a direct protein-protein interaction, facilitating an alternative, poly(A)-independent, closed-loop structure (14, 62). This activity might enable translational activation of germ-cell mRNAs without prior polyadenylation. Nevertheless, a subset of DAZL target mRNAs also contain cytoplasmic polyadenylation elements (CPEs) (11, 54, 64), suggesting the possibility of synergistic or sequential regulation by DAZL and CPE-binding protein (CPEB), which promotes polyadenylation (57, 64). Indeed, we have previously shown that DAZL can further augment translation of polyadenylated mRNA (14).

Interestingly, a study in zebrafish suggested that DAZL may play a direct role in cytoplasmic polyadenylation (66). Ectopic expression of DAZL within somatic cells resulted in an increase in reporter mRNA poly(A)-tail length compared with cells devoid of DAZL or those expressing a DAZL mutant deficient in RNA binding (66). This led the authors to conclude that DAZL might be able to induce cytoplasmic polyadenylation, although this proposed activity was not directly examined. This study raises the possibility that DAZL may activate translation in both poly(A)-independent and poly(A)-dependent manners.

In our present study we directly investigate the ability of DAZL to promote cytoplasmic polyadenylation. Using a tether-function approach in *X. laevis* oocytes we find that DAZL does not induce polyadenylation but confirm by RNA immunoprecipitation that DAZL can recruit PABP directly to unadenylated mRNAs to activate translation. Importantly, we reveal a previously undescribed activity of DAZL in inhibiting mRNA deadenylation. We find that this activity is dependent upon the interaction of DAZL with PABP but, surprisingly, independent of the role of the DAZL-PABP complex in activating initiation of translation and indeed independent of translation *per se*, showing that these events are mechanistically separable. Inhibition of deadenylation is not mediated through the C-terminal region of PABP, which binds deadenylases (6), but does require the poly(A)-binding activity of PABP, suggesting a model in which DAZL aids the recruitment and/or retention of PABP on the poly(A) tail, sterically occluding the action of deadenylases. Our findings shed light on the poorly understood relationship between deadenylation and translation by revealing that these activities are uncoupled for DAZL-PABP complexes.

## RESULTS

### DAZL does not require mRNA polyadenylation to activate translation

Previous work led to a model for DAZL-mediated activation of translation in which DAZL recruits PABP directly to mRNA, effectively bypassing the requirement for the poly(A) tail for PABP recruitment (14). However, subsequent work in zebrafish suggested that DAZL may induce mRNA polyadenylation (66), raising the possibility that it can stimulate translation via both poly(A)-dependent and independent mechanisms. We therefore sought to clarify its mechanism(s) of action by directly assessing whether the DAZL-PABP interaction is indeed sufficient to recruit PABP in the absence of a poly(A) tail and whether DAZL can mediate cytoplasmic polyadenylation. In this study we focused on murine DAZL since this is currently the most extensively analysed homologue at the molecular level (14, 62, 64) and primarily adopted an MS2-based tether-function approach in *X. laevis* oocytes (14, 25) since tethering avoids the inclusion of other *cis*-acting control elements that may be present within natural 3’UTRs and that might confound mechanistic analysis. Furthermore, both regulated changes in poly(A)-tail length and DAZL function have been abundantly studied in these germ cells (7).

During *X. laevis* oocyte maturation, mRNAs containing the requisite control elements undergo cytoplasmic polyadenylation coincident with their translational activation (74). For instance, a microinjected, *in vitro* transcribed, unadenylated luciferase reporter mRNA containing the cytoplasmic polyadenylation elements (CPEs) and hexanucleotide sequence (Hex) from the cyclin B 3’UTR (Luc-MS2-cycB1, Fig. S1 [1]) (3, 25) becomes polyadenylated upon progesterone-induced oocyte maturation (Fig. 1 *A*, lanes 9 & 11). This was analysed with a ligation- and RT-PCR-dependent poly(A)-tail (PAT) assay (9) and verified by treatment with oligo(dT) and RNase H treatment (Fig. S2*A*). As expected, polyadenylation correlated with an increase in luciferase activity normalised to β-galactosidase, expressed from an internal control mRNA lacking polyadenylation sequences (Fig. S2*B*; Fig. S1 [16]). These data show that oocyte maturation was occurring successfully in our system.

**Fig. 1.**
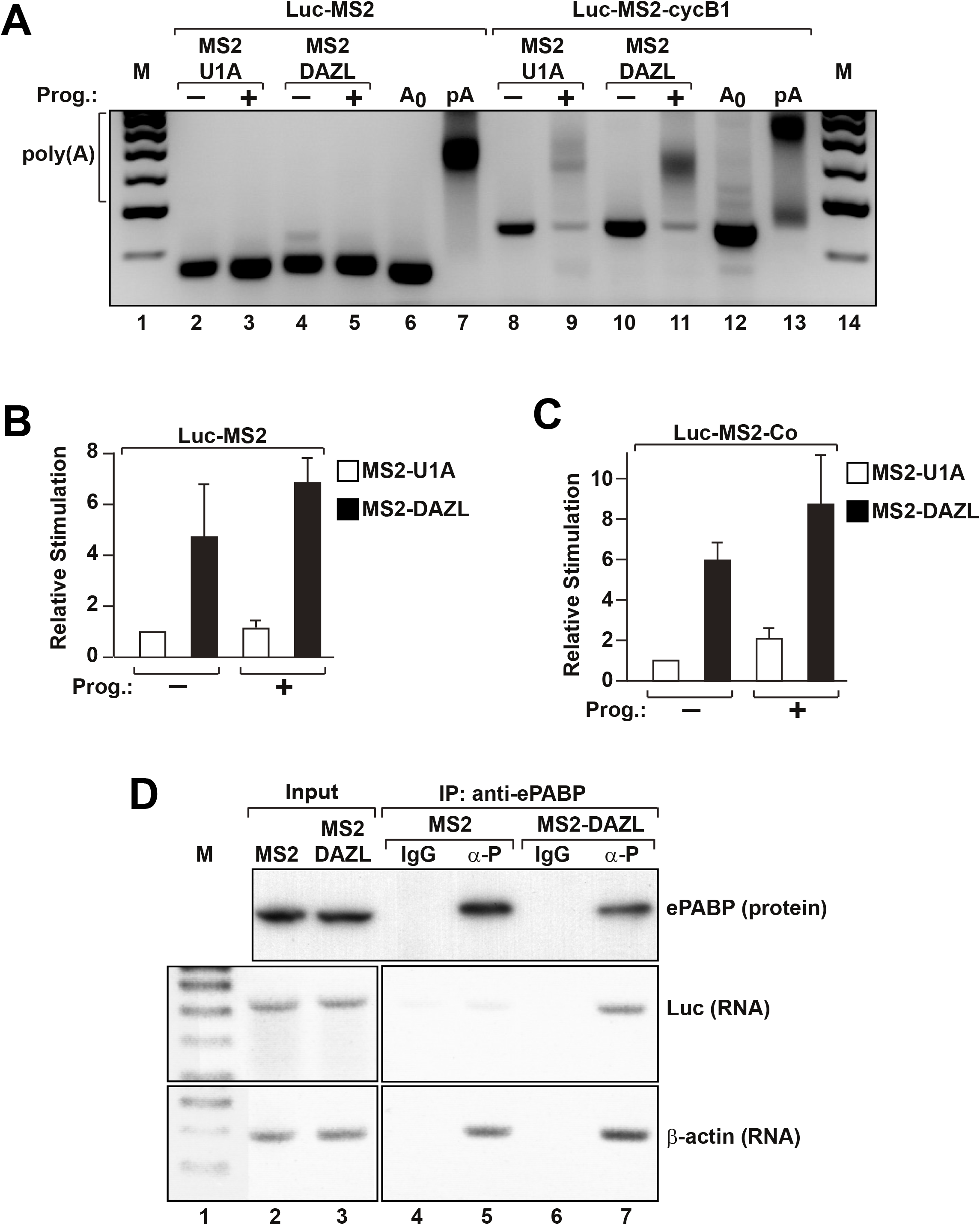
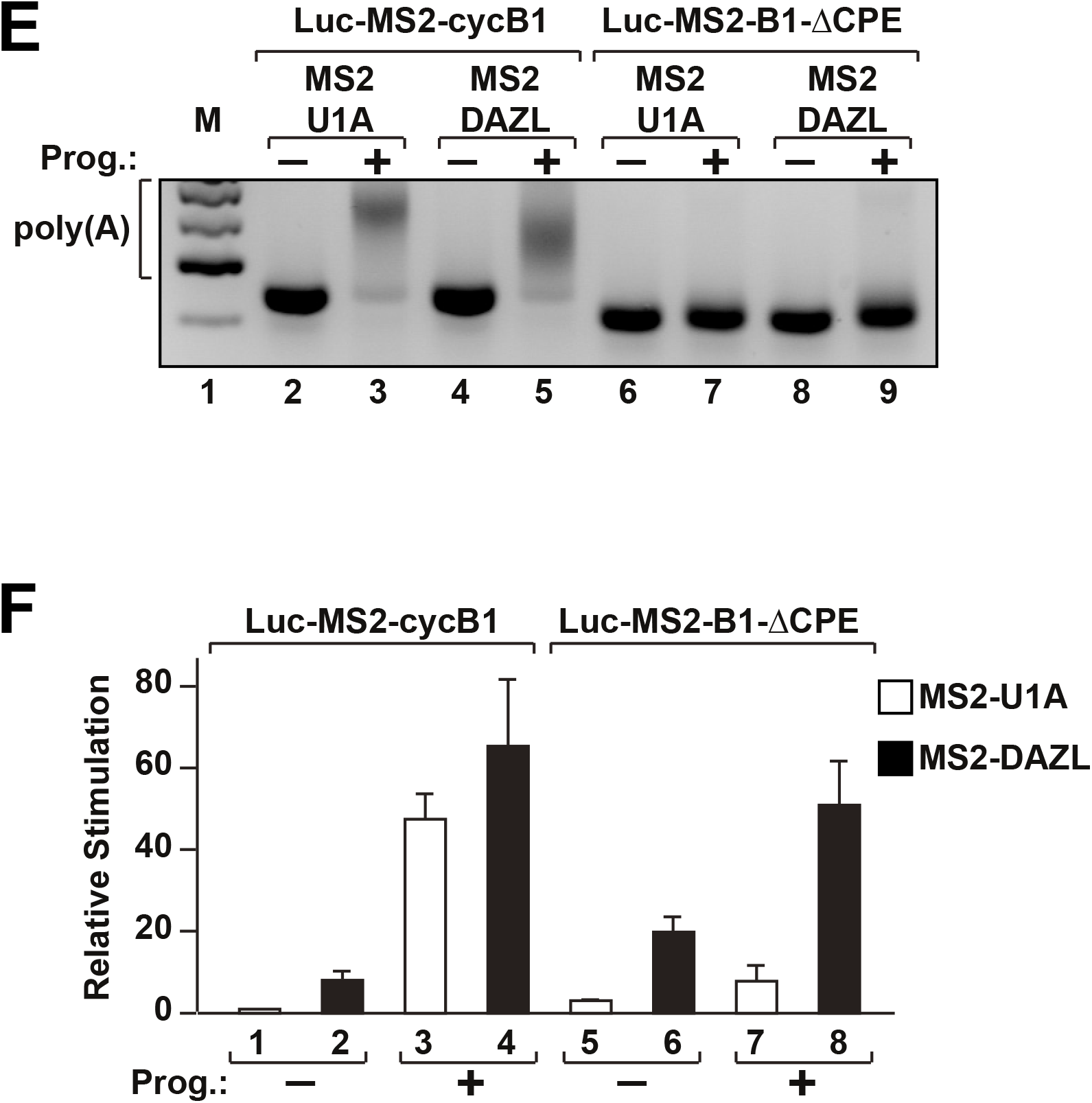
Polyadenylation is not induced by DAZL, nor is it required for DAZL-mediated translational activation. ***(A)*** Oocytes expressing MS2-U1A or MS2-DAZL were co-injected with either Luc-MS2 or Luc-MS2-cycB1 and CSFV-β-gal (internal control) mRNAs and treated with progesterone (Prog.) (+) or left untreated (-). mRNA polyadenylation was analysed by PAT assay. M, DNA size markers; poly(A), high molecular weight bands indicative of polyadenylation; A_0_ & pA, polyadenylation controls: PAT assays of unadenylated and *in vitro* polyadenylated reporter mRNA. ***(B)*** Effects of MS2-U1A and MS2-DAZL on translation of Luc-MS2 (progesterone-treated or untreated; see (*A*)) were measured by luciferase assay normalised to β-galactosidase activity. Effects on translation of relevant samples from (*A*) were measured by luciferase assay normalised to β-galactosidase activity. Translational stimulation relative to Luc-MS2 in untreated MS2-U1A-expressing cells (value set to 1) is plotted. Error bars represent standard error of the mean (SEM); *n*=3. (***C***) Effects on translation as in (*B*) but with Luc-MS2-Co were measured as in (*B*) but with Luc-MS2-Co, MS2-U1A (-) set to 1; *n*=3. ***(D)*** Oocytes expressing MS2 or MS2-DAZL were injected with ApG-capped Luc-MS2. ePABP was immunoprecipitated (α-P) from lysates (control: non-specific rabbit IgG) and detected by immunoblotting. Input represents 10%. The presence of co-purified luciferase or endogenous β-actin mRNAs was assessed by RT-PCR. Input represents 24%. ***(E)*** Oocytes expressing MS2-U1A or MS2-DAZL were co-injected with either Luc-MS2-cycB1 or Luc-MS2-B1-ΔCPE and CSFV-β-gal mRNAs, progesterone treated (+) or not (-), and analysed as in (*A*). ***(F)*** Effects of MS2-U1A and MS2-DAZL on translation of Luc-MS2-cycB1 and Luc-MS2-B1-ΔHex (progesterone-treated or untreated; see *(E))* were analysed as in (*B*) with Luc-MS2-cycB1, MS2-U1A (-) set to 1; *n*=3.

We previously showed that translation of an unadenylated reporter mRNA (Luc-MS2; Fig. S1 [4]) can also be activated by tethered DAZL (14, 62), however, in those studies the polyadenylation status of Luc-MS2 had not been examined after micro-injection and was therefore analysed here. We observed no polyadenylation in fully grown (stage VI) oocytes expressing either the negative control MS2-U1A (a translationally inert RNA-binding protein) or MS2-DAZL pre- or post-maturation (Fig. 1*A*, lanes 2-5), consistent with the absence of polyadenylation elements in Luc-MS2. Nevertheless, MS2-DAZL robustly stimulated translation (Fig. 1 *B*), strongly suggesting that this is a poly(A)-independent activity. Despite the absence of key polyadenylation elements in Luc-MS2, we cannot entirely exclude sub-detection polyadenylation by an unknown mechanism; for instance, Hex-independent cytoplasmic polyadenylation has been described in *Drosophila melanogaster* (12). Since poly(A) polymerase activity is dependent on the presence of a 3’ hydroxyl group on the RNA we added 3’ deoxyadenosine (cordycepin) to the Luc-MS2 3’ end (Luc-MS2-Co; Fig. S1 [5]) prior to microinjection to chemically prohibit nucleotidyl terminal transfer. Cordycepin incorporation was confirmed using an RNA ligation-dependent RT-PCR reaction (Fig. S2*C*) but was found not to prevent DAZL-dependent translational activation (Fig. 1*C*; Fig. S2*D*).

### DAZL recruits PABP to unadenylated mRNA

Having shown that cytoplasmic polyadenylation is not an *a priori* requirement for translational stimulation by DAZL, we sought to confirm whether DAZL can recruit PABP poly(A)-independently via a direct protein-protein interaction as previously proposed (14). RNA co-immunoprecipitations of a tethered unadenylated reporter mRNA (ApG-Luc-MS2; Fig. S1 [4]) were performed using antibodies against embryonic PABP (ePABP), the predominant PABP in oocytes (7) (Fig. 1 *D*). The mRNA was non-physiologically (ApppG) capped to avoid possible pull-down via an ePABP-eIF4G (77), rather than through an ePABP-DAZL, interaction. Critically, ePABP failed to associate with Luc-MS2 unless MS2-DAZL, rather than MS2 alone, was present (Fig. 1*D*, lanes 4 & 6). Endogenous polyadenylated β-actin mRNA, which binds ePABP directly via its poly(A) tail, was a positive control (Fig. 1*D*, lower panels). Thus, MS2-DAZL suffices to mediate the association of ePABP with the reporter mRNA, providing further support for a mechanism through which DAZL activates translation initiation by directly recruiting PABP.

### DAZL does not induce polyadenylation in *X. laevis* oocytes

Our data reveal that DAZL-mediated translational stimulation occurs via direct protein-mediated recruitment of PABP and is independent of cytoplasmic polyadenylation, but do not directly address whether DAZL might promote polyadenylation in a manner similar to CPEB. Thus the CPEs, but not the Hex, were deleted from Luc-MS2-cycB1 (Luc-MS2-B1-ΔCPE; Fig. S1 [2]) abrogating polyadenylation (Fig. 1*E*, lanes 6 & 7) in mature control (MS2-U1A expressing) oocytes, as previously described (67). Importantly, no appreciable polyadenylation of Luc-MS2-B1-ΔCPE occurred in the presence of tethered DAZL (lanes 8 & 9), demonstrating that it does not induce polyadenylation. (Although a minimal increase in product size appears to be consistently present upon maturation (lanes 7 & 9), this is not DAZL-specific.) These data are consistent with experiments using other reporter mRNAs, either lacking the Hex sequence or the entire cyclin B1 3’ UTR (Luc-MS2-B1-ΔHex & Luc-MS2; Fig. S1 [3] & [4]) also showing no detectable DAZL-mediated mRNA polyadenylation (Fig. S2*E*, lanes 11-18; Fig. 1*A*, lanes 2-5). DAZL-mediated translational stimulation of Luc-MS2-B1-ΔCPE was robust (Fig. 1*F*) although an increase in basal translation was observed (Fig. 1*F*, bars 1-2 *vs* 5-6) consistent with relief of CPEB-mediated translational repression in immature oocytes (3). Similarly, translational stimulation of Luc-MS2-B1-ΔHex was efficient (Fig. S2*F*), consistent with our finding that polyadenylation is not required for DAZL-mediated translational stimulation.

### DAZL inhibits mRNA deadenylation in *X. laevis* oocytes

Since we found no evidence of mRNA polyadenylation by DAZL, we hypothesised that the apparent DAZL-dependent poly(A) tail extension observed in zebrafish cells (66) may reflect an effect of DAZL on deadenylation rather than polyadenylation, since both processes contribute to poly(A)-tail length. Upon *X. laevis* oocyte maturation, mRNAs with long poly(A) tails that lack *cis*-acting cytoplasmic polyadenylation elements undergo so-called ‘default deadenylation’ (20). Fig. S3*A* shows that this process leads to the gradual and complete removal of the poly(A) tail from an *in vitro* polyadenylated reporter mRNA (Luc-MS2-pA; Fig. S1 [4]) micro-injected into the cytoplasm of oocytes that were subsequently treated with progesterone. Strikingly, this process is strongly inhibited in oocytes expressing MS2-DAZL (Fig. 2*A*), demonstrating a previously undescribed activity of DAZL in regulating mRNA deadenylation, which is conserved between mouse and frog (Fig. S3*B*). This activity is abolished by deletion of the MS2-binding sites from the reporter mRNA (Luc-ΔMS2-pA; Fig. S1 [6]) demonstrating that DAZL must be bound to mRNA to regulate poly(A) tail length (Fig. 2*B*).

**Fig. 2.**
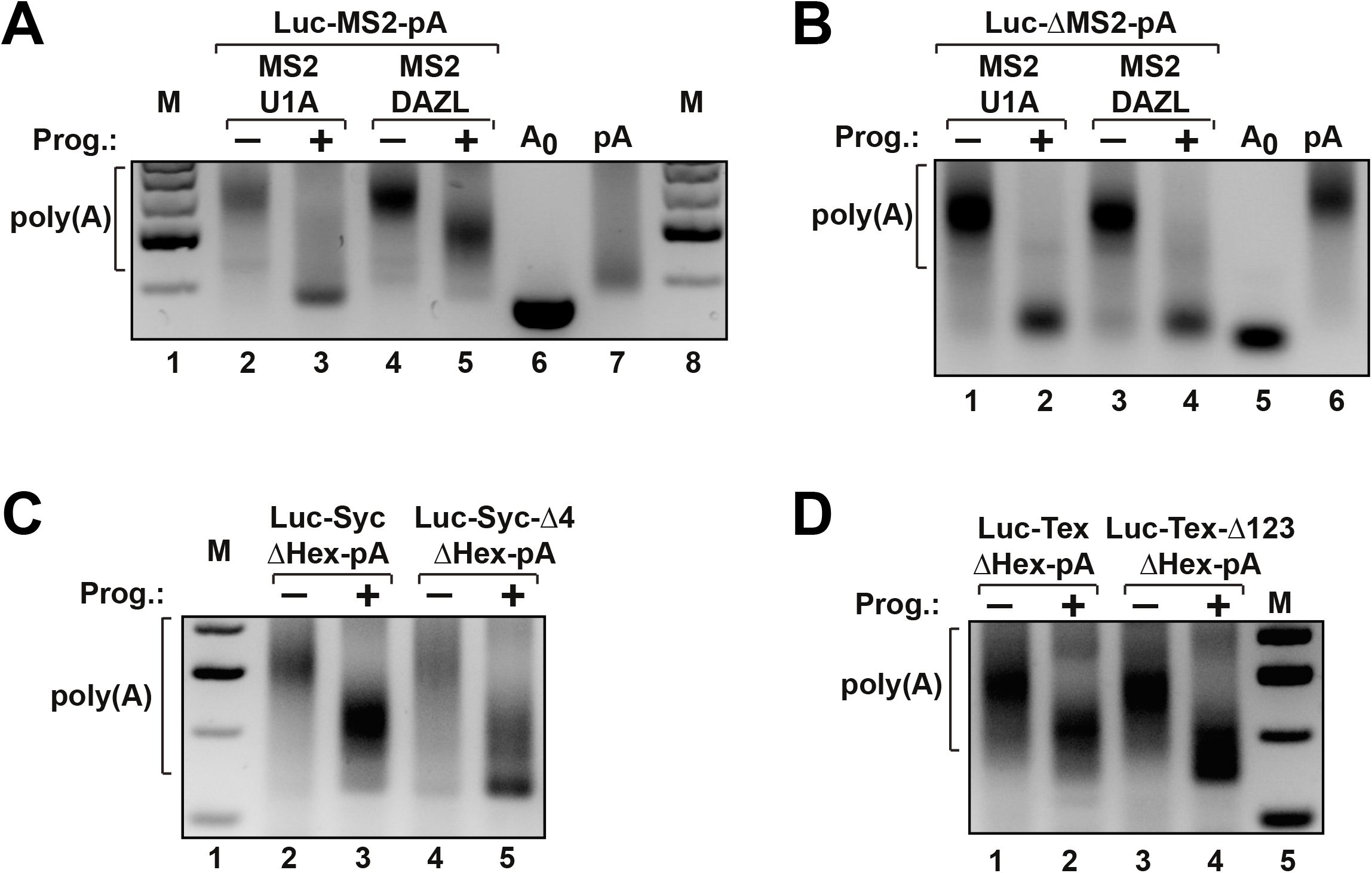
DAZL inhibits mRNA deadenylation. ***(A)*** Oocytes expressing MS2-U1A or MS2-DAZL were injected with Luc-MS2-pA, progesterone-treated (+) or not (-), and analysed as in Fig. 1*A*. ***(B)*** Oocytes expressing MS2-U1A or MS2-DAZL were injected with Luc-ΔMS2-pA and analysed as in Fig. 1*A*. ***(C)*** Oocytes were either co-injected with Luc-Syc-ΔHex-pA and Luc-MS2-pA or with Luc-Syc-Δ4-ΔHex-pA and analysed as in Fig. 1*A*. See also Fig. S3E. ***(D)*** Oocytes were injected with Luc-Tex-ΔHex-pA or Luc-Tex-Δ123-ΔHex-pA and analysed as in Fig. 1*A*.

To investigate whether murine DAZL can similarly inhibit deadenylation of validated target mRNAs, we used reporter mRNAs with either the murine *Sycp3* or *Tex19.1* 3’UTRs, previously shown to be responsive to DAZL-mediated translational stimulation (11, 54, 64) (Fig. S1 [7] & [10]). Since these mRNAs contain elements that promote maturation-dependent polyadenylation (Fig. S3*C* & S4*D*, lanes 2 & 3), we mutated the Hex to abolish this property (Fig. S1 [8] & [11]; Fig. S3*C* & S4*D*, lanes 4 & 5). The resulting reporter mRNAs were polyadenylated *in vitro* (Luc-Syc-ΔHex-pA & Luc-Tex-ΔHex-pA) and micro-injected into *X. laevis* oocytes. During progesterone treatment, deadenylation was inefficient compared with co-injected Luc-MS2-pA (Fig. 2*C*, lanes 2 & 3 and Fig. 2*D*, lanes 1 & 2 vs Fig. S3*E*).

We posited that this was due to the *Sycp3* and *Tex19.1* 3’UTRs being bound by endogenous *X. laevis* DAZL, which is 83% identical within the RRM domain with murine DAZL (likely recognising similar RNA motifs) and shares its ability to inhibit deadenylation (Fig. S3*B*). To test this, we mutated putative DAZL-binding sites within the 3’UTRs, including mutations previously shown to abrogate DAZL binding (Table S1; Fig. S1 [9] & [12]) (54, 64). Deadenylation of the mutated 3’UTRs was considerably more efficient than that of the wildtypes, indicating an ability of non-tethered DAZL to regulate the poly(A)-tail length of its natural target mRNAs.

### Inhibition of deadenylation requires the PABP-binding region of DAZL

To explore the mechanism responsible for DAZL-mediated impairment of deadenylation a possible role of its partner protein PABP was explored for several reasons: PABP has been reported to associate with and regulate the activity of various deadenylase enzyme complexes (6, 10, 19, 49, 72, 81) and its regulatory role in translation initiation might influence poly(A) tail protection since mRNA stability, of which deadenylation is the rate limiting step, and translation are intricately linked (52), although the mechanistic basis of this is has yet to be elucidated.

The minimal DAZL region required for PABP binding was previously mapped to amino acids 99-166, partially overlapping the RRM (14) (Fig. 3*A*), although amino acids outside this region *(e.g.* the DAZ motif) also influence PABP binding (14, 78). Deletion of this region entirely abolished the ability of DAZL to inhibit deadenylation (Fig. 3*B*) despite efficient expression of mutant DAZL in oocytes (Fig. S4*A*), whereas deletion of both the N-terminal 32 and C-terminal 108 amino acids (DAZL 33-190), outside the PABP-binding region, had no effect on poly(A) stabilisation by DAZL (Fig. 3*A*, *C*). These findings suggested that this activity could be mediated by a DAZL-PABP interaction, predicting that direct tethering of PABP or of another PABP-binding protein would likewise impede deadenylation. Indeed, direct tethering of PABP1 (Fig. 3*D*) and tethering of the PABP-binding region of a PABP-dependent viral translational activator unrelated to DAZL, ICP27 (62), recapitulated the effect of DAZL (Fig. 3*E*). Together these data strongly indicate that DAZL regulates poly(A) tail length via its interaction with PABP.

**Fig. 3.**
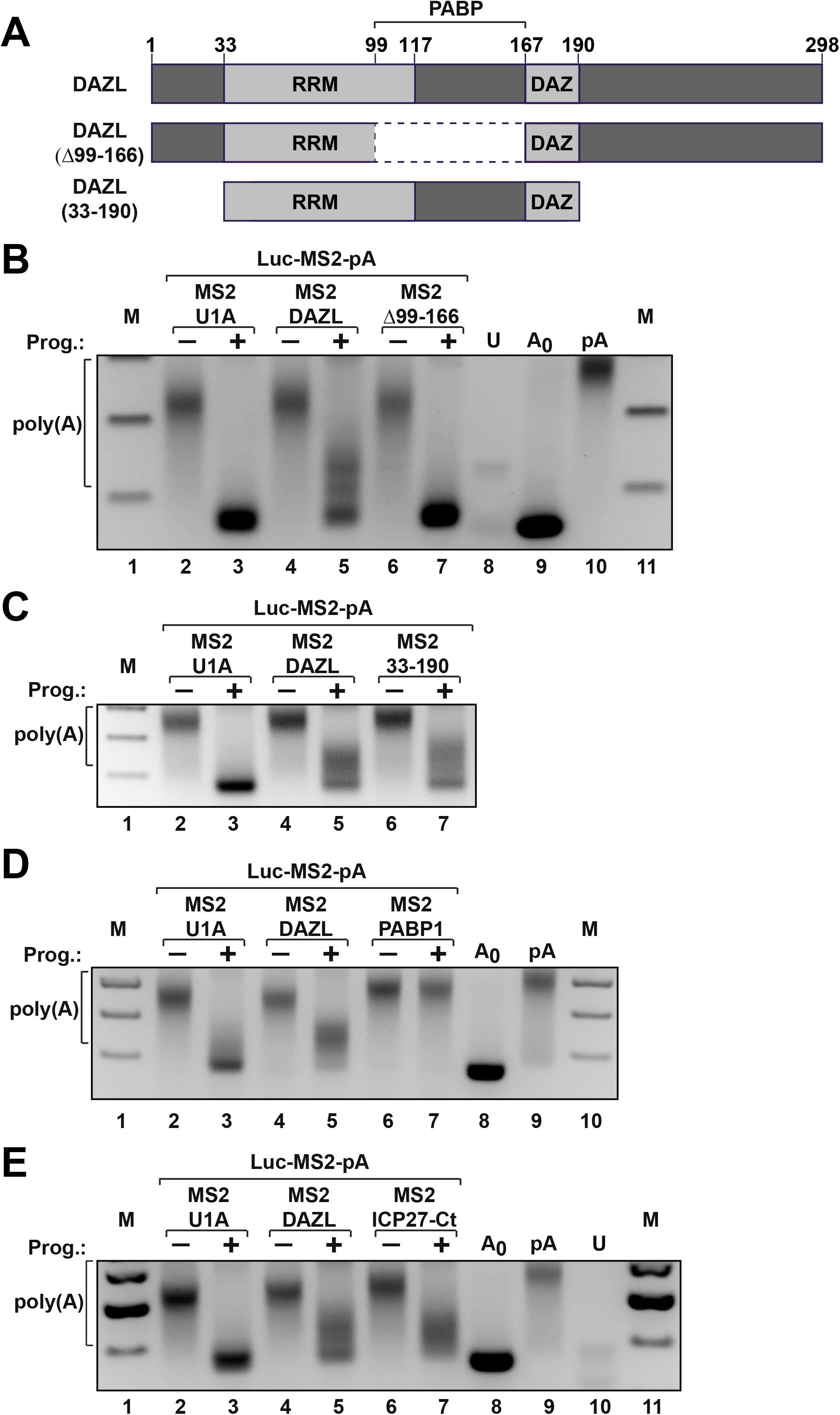
Inhibition of deadenylation requires PABP. **(*A*)** Top: DAZL domain organisation showing the location of the RRM and DAZ motifs and the minimal region required for PABP binding (14). Middle and Bottom: Protein regions retained in the DAZL mutants used in this study. Dashed lines indicate absent regions; numbers indicate positions of amino acid residues. **(*B*)** Oocytes expressing MS2-U1A, MS2-DAZL or MS2-DAZL (Δ99-166) were co-injected with Luc-MS2-pA, treated and analysed as in Fig. 1*A*. **(*C*)** Oocytes expressing MS2-U1A, MS2-DAZL or MS2-DAZL (33–190) were co-injected with Luc-MS2-pA, treated and analysed as in Fig. 1*A*. **(*D*)** Oocytes expressing MS2-U1A, MS2-DAZL or MS2-PABP1 were co-injected with Luc-MS2-pA, treated and analysed as in Fig. 1*A*. ***(E)*** Oocytes expressing MS2-U1A, MS2-DAZL or MS2-ICP27-Ct (amino acids 201-512) were co-injected with Luc-MS2-pA, treated and analysed as in Fig. 1*A*.

### Inhibition of deadenylation by DAZL does not require translational activation

We have shown that PABP recruitment to an mRNA by DAZL, ICP27 or direct tethering inhibits default deadenylation. However, the same complexes also activate translation initiation in immature (14, 25, 38, 62) and mature oocytes (Fig. 1*B* & Fig. S4*B*). Since it is increasingly apparent that translational efficiency can be an important determinant of deadenylation and mRNA stability (46, 51, 52), this raised the possibility that PABP could be acting indirectly. Therefore, we investigated in several ways whether DAZL-mediated deadenylation inhibition might be a consequence of enhanced ribosome association with Luc-MS2-pA. Initially, we tethered the PABP-independent, mRNA-specific translational activator, stem loop-binding protein (SLBP) (23) to Luc-MS2-pA and found that it failed to inhibit deadenylation (Fig. 4*A*, lanes 6 & 7), suggesting that PABP recruitment rather than translational activation *per se* (Fig. S4*C*) was required for inhibition of deadenylation. In a complementary approach we tethered DAZL to the CSFV-Luc-MS2-pA reporter mRNA (Fig. S1 [14]), translation initiation of which is driven by the classical swine fever virus internal ribosome entry site (CSFV IRES). CSFV IRES-dependent translation is efficient but independent of the eIF4G-containing cap-binding complex (50) and is not enhanced by PABP or DAZL (62). Nevertheless, DAZL efficiently inhibits deadenylation of this reporter mRNA (Fig. 4*B*), indicating that translational stimulation is not a prerequisite for its activity. In order to examine whether basal translation is required for DAZL-mediated retardation of deadenylation, we tethered DAZL to two mRNAs bearing an ApppG-cap structure (ApG-Luc-MS2-pA; Fig. S1 [4]), one of which additionally contained a stable hairpin in the 5’UTR (ApG-HP-Luc-MS2-pA; Fig. S1 [13]). The ApppG cap is not recognised by the cap-binding complex leading to a reduction of translational efficiency by two orders of magnitude (21) (Fig. S4*D*). Additional introduction of a −50kcal/mol hairpin further reduced reporter gene expression to undetectable levels (2) (Fig. S4*D*). Nevertheless deadenylation of both constructs was efficiently inhibited (Fig. 4*C*; Fig. S4*E*), clearly indicating, alongside our other data, that this activity is independent of translation levels and any effects of DAZL thereupon.

**Fig. 4.**
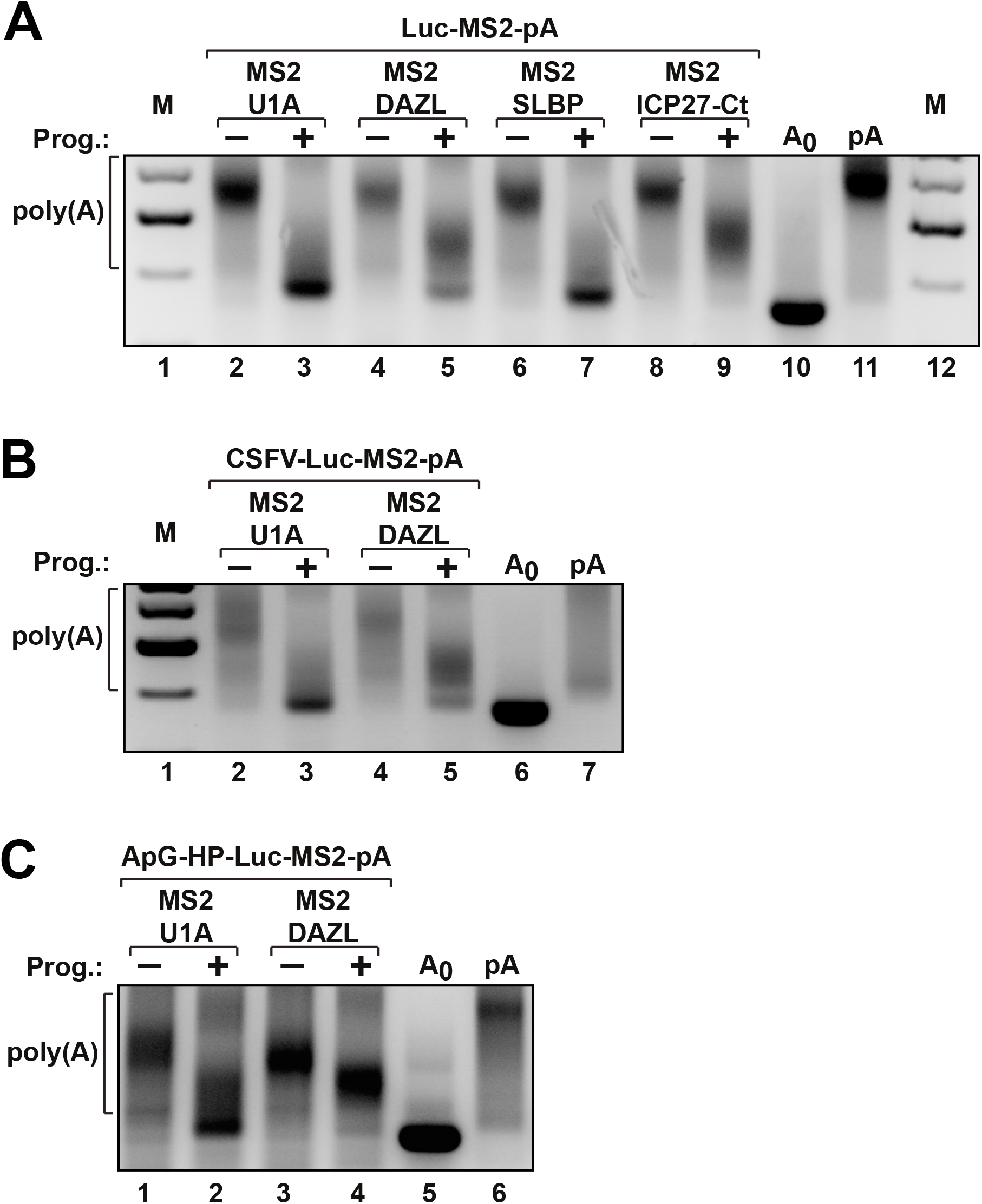
DAZL-mediated inhibition of deadenylation is independent of translational activation. **(*A*)** Oocytes expressing MS2-U1A, MS2-DAZL, MS2-ICP27-Ct or MS2-SLBP were co-injected with Luc-MS2-pA, treated and analysed as in Fig. 1*A*. **(*B*)** Oocytes expressing MS2-U1A or MS2-DAZL were co-injected with CSFV-Luc-MS2-pA, treated and analysed as in Fig. 1*A*. **(*C*)** Oocytes expressing MS2-U1A or MS2-DAZL were co-injected with ApG-HP-Luc-MS2-pA, treated and analysed as in Fig. 1*A*.

### Inhibition of deadenylation requires functional PABP1 RRMs

Having established that recruitment of PABP rather than its effect on translation is key to the poly(A)-protective effect of DAZL, we analysed the activity of directly tethered PABP1 in deadenylation inhibition (Fig. 3*E*) to gain further insight into the underlying mechanism.

PABP has been proposed to link translation and deadenylation through its role in forming an ‘end-to-end’ ribonucleoprotein complex via its interaction with eIF4G, which enhances translation, by protecting the mRNA from deadenylases either sterically or as a result of competition between translation factors and deadenylases for binding the C-terminal PABC domain (40). Methionine 161 in PABP1 RRM2 is critical for its interaction with eIF4G, and translational activation, but not for PABP-poly(A) binding (26, 32). Intriguingly, substitution of this amino acid did not affect the ability of tethered PABP1 (M161V) to inhibit deadenylation (Fig. *5A*) despite dramatically reducing its ability to activate translation (Fig. S5*A*) due to a compromised ability to promote end-to-end complex formation. This result is consistent with the independence of DAZL-mediated inhibition of deadenylation on active translation. We confirmed similar separation of translation and poly(A)-tail stabilisation for PABP1 using a non-physiologically capped mRNA lacking an open reading frame (ΔORF-MS2-pA; Fig. S1 [15]) that is not a substrate for translation. Deadenylation of this mRNAs is inhibited by tethered PABP (Fig. 5*B*).

**Fig. 5.**
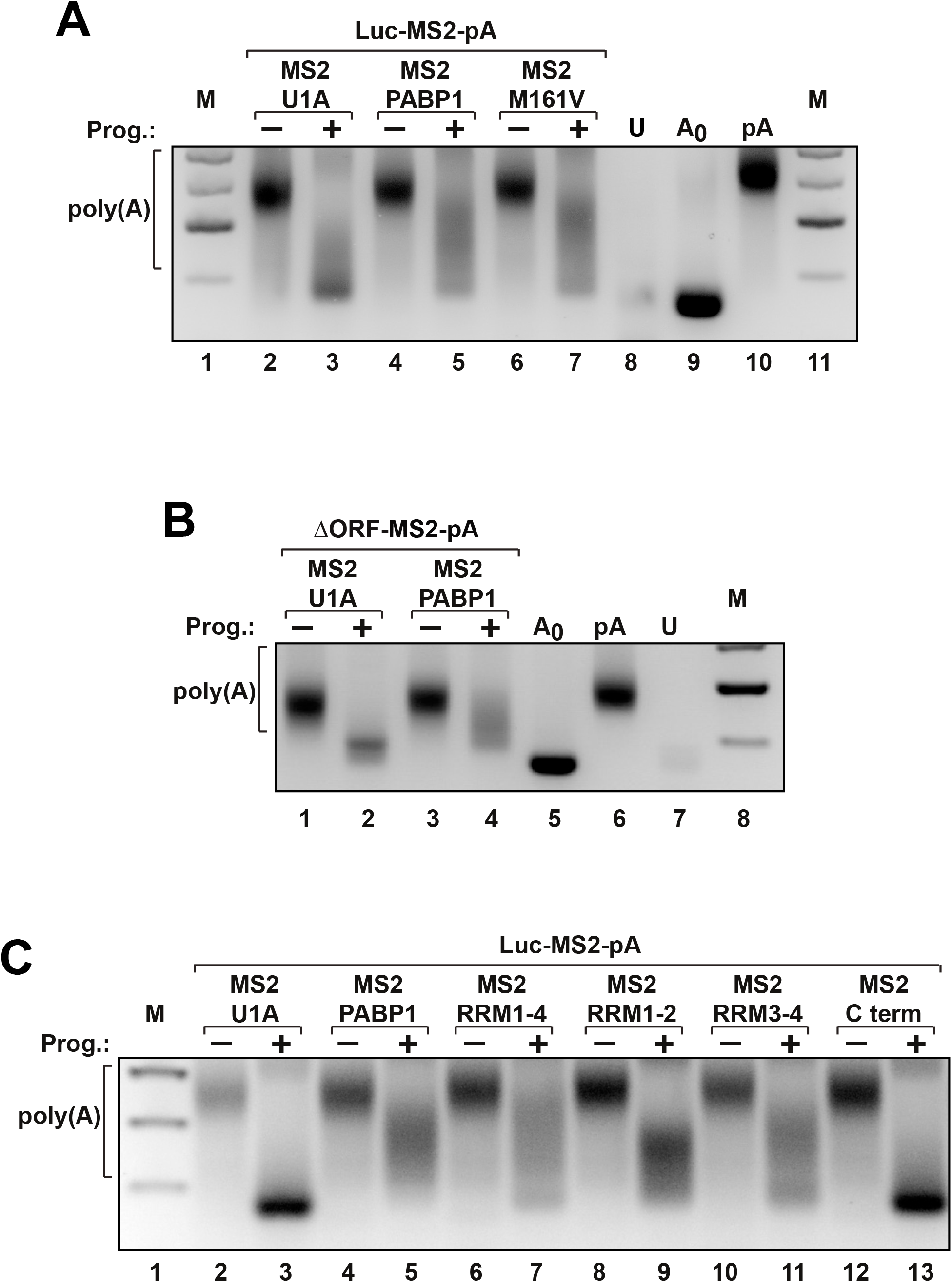
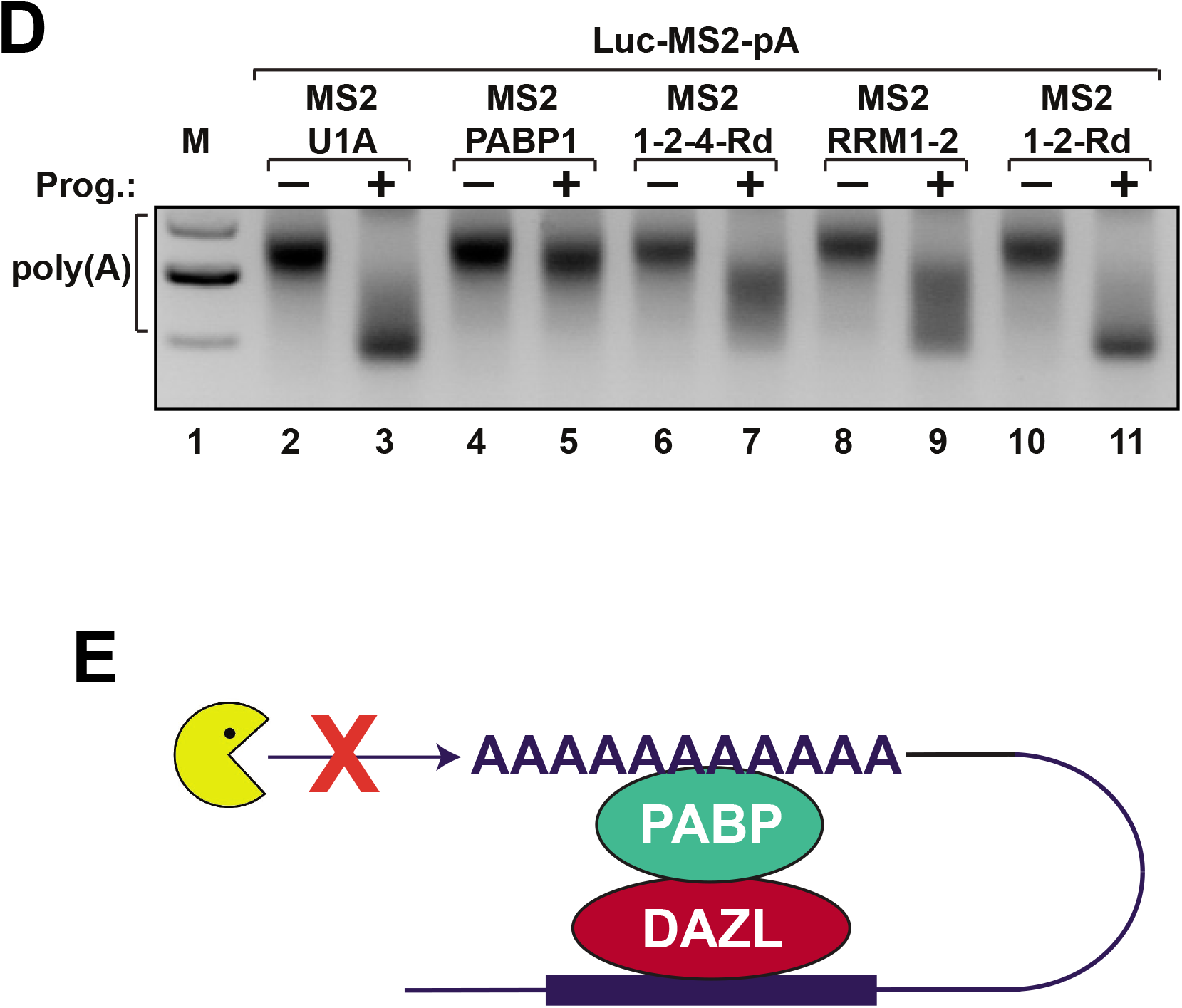
The PABP RRMs must bind RNA to impede deadenylation. **(*A*)** Oocytes expressing MS2-U1A, MS2-PABP1 or MS2-PABP1 (M161V) were co-injected with Luc-MS2-pA, treated and analysed as in Fig. 1*A*. **(*B*)** Oocytes expressing MS2-U1A or MS2-PABP1 were co-injected with ApG-ΔORF-MS2-pA and analysed as in Fig. 1*A*. **(*C*)** Oocytes expressing MS2-U1A, MS2-PABP1, MS2-PABP1 (RRM1-4), MS2-PABP1 (RRM1-2), MS2-PABP1 (RRM3-4) or MS2-PABP1 (C term.) were co-injected with Luc-MS2-pA, treated and analysed as in Fig. 1*A*. **(*D*)** Oocytes expressing MS2-U1A, MS2-PABP1, MS2-PABP1 (RRM1-2-4Rd), MS2-PABP1 (RRM1-2) or MS2-PABP1 (RRM1-2Rd) were co-injected with Luc-MS2-pA, treated and analysed as in Fig. 1*A*. Mutation of all four RRMs resulted in loss of PABP1 translation activity (data not shown) indicating non-specific effects and so this construct was not included in our study. **(*E*)** A model for DAZL-mediated inhibition of deadenylation. DAZL recruits PABP to the 3’UTR of specific mRNAs thereby stabilising the interaction of PABP with the poly(A) tail and impeding the access or inhibiting the activity of deadenylases, possibly serving to maintain some mRNAs in a translationally active state once they have been polyadenylated by other factors.

As neither the ability of PABP1 to stimulate translation or participate in end-to-end complex formation were required for its ability to impede deadenylation we exploited the advantages of the tether-function system to analyse the contribution of individual PABP1 domains (25). PABP1 comprises four nonidentical RRMs with different RNA-binding specificities and a C-terminal region (PABC) that does not bind RNA but interacts with numerous proteins, including several deadenylases (7) (Fig. S5*B*). We tethered the combinations of domains shown in Fig. S5*B* separately to Luc-MS2-pA revealing that the RRMs but not the C-terminal region are sufficient to inhibit deadenylation (Fig. 5*C*), indicating that PABC-deadenylase interactions do not play a role in this activity. Rather, despite PABP1 being tethered to the reporter mRNA via MS2, the RNA-binding function of PABP1 appeared to be required. This was confirmed by introducing point mutations that specifically abolish the RNA-binding capacity of the individual RRMs without affecting other known activities of PABP1 (25). Mutating RRM 1 & 2 abolished the protective effect of this region tethered in isolation (Fig. 5*D*, lanes 8-11) but did not impair the ability of this region to stimulate translation (25), further underlining the mechanistic independence of these PABP-mediated events. Similarly, mutation of RRM 1, 2 & 4 drastically reduced the poly(A)-protective effect of full-length PABP1 (Fig. 5*D*, lanes 4-7). We conclude that recruited PABP1 must bind the poly(A) tail in order to ensure its stabilisation, leading to a model, illustrated in Fig. 5*E*, whereby DAZL-recruited PABP also interacts with the poly(A) tail and in so doing sterically occludes the action of deadenylases.

## DISCUSSION

In this study we have examined the role of DAZL in regulating poly(A)-tail length. Using the tether-function assay we investigated the possibility that mammalian DAZL may induce cytoplasmic mRNA polyadenylation but found no evidence to support this (Fig. 1 *E*). Rather, our study revealed that binding of DAZL to an mRNA, either by tethering or via natural binding motifs, inhibits cytoplasmic mRNA deadenylation (Fig. 2–4) via recruitment of PABP (Fig. 3*D* & 5) and that this activity requires the RNA-binding capacity of PABP (Fig. 5), leading to the model depicted in Fig. 5*E*, where enhanced PABP occupancy of the poly(A) tail occludes deadenylase access. Furthermore, inhibition of deadenylation by the DAZL-PABP complex does not require the RNA to be translationally competent and is therefore independent of its role in mRNA-specific translational activation (Fig. *4C* & Fig. 5*B*)

In our system, mammalian DAZL does not induce polyadenylation during oocyte maturation on mRNAs lacking a CPE but retaining a functional Hex, thought to be required for all cytoplasmic polyadenylation in vertebrates (76) (Fig. 1 *E*). Furthermore, we did not observe increased polyadenylation of a CPE-containing mRNA in the presence of DAZL (Fig. 1*A, E* & Fig. S2*E*), despite a report that DAZL and CPEB regulate mRNA translation synergistically (64), suggesting that such synergism does not arise from increased polyadenylation. Although a direct role in polyadenylation was not determined in zebrafish, our present data do not rule out that zebrafish DAZL may behave differently in this respect or indeed that the activity of (mammalian) DAZL may be altered in zebrafish cells, but eliminates DAZL-mediated polyadenylation as a requirement for mRNA-specific translational activation by mammalian and another non-mammalian vertebrate DAZL (14, 54, 55). Consistent with its ability to recruit PABP to unadenylated mRNA (Fig. 1 *D*), we confirmed poly(A)-independent translational activation by DAZL with different reporter constructs, including those on which polyadenylation should be largely blocked by 3’ cordycepin (Fig. 1). Cordycepin is likely to remain stably incorporated during oocyte maturation, preventing polyadenylation throughout this process, since it is not a substrate for poly(A)-specific ribonuclease (PARN) (1), the deadenylase responsible for the default deadenylation process in *X. laevis* oocytes (37). Furthermore, direct recruitment of PABP by DAZL is also consistent with our previous findings that DAZL, like PABP, stimulates poliovirus (PV) IRES-dependent but not CSFV IRES-dependent initiation of translation (62).

Our discovery that DAZL retards maturation-dependent default deadenylation (Fig. 2) suggests that it inhibits the activity of PARN, since this is the main default deadenylase (15, 37). This is consistent with the requirement for PABP for DAZL activity (Fig. 3) since *X. laevis* maturation-specific deadenylation has been shown to be antagonised by PABP1 overexpression (79) but accelerated by ePABP depletion (71) and mammalian PARN is inhibited by PABP (36). However, despite reports that PARN activity is stimulated by a 5’ cap *in vitro* (16), we observed efficient deadenylation of our non-physiologically capped reporter mRNAs in the absence of DAZL. We suggest that this is due to the cap requirement for PARN activity not being absolute (16, 44) allowing deadenylation during the time course of our assays, but it is also possible that the cap requirement is reduced *in vivo*. Although we suggest that PARN is the likely target of inhibition by the DAZL-PABP complex in our assays, we do not rule out that it may also impede the action of other deadenylases present in *X. laevis* oocytes (22). However, the relationship between PABP and deadenylation is complex with PABP both stimulating and suppressing the activities of different deadenylases (6, 72, 81). It is noteworthy that the C-terminal region of PABP, which mediates interactions with several major deadenylase complexes (poly(A) nuclease [PAN]2-PAN3 and carbon catabolite repressor protein [CCR]4-negative on TATA [NOT]) (6) is dispensable for DAZL-mediated inhibition of deadenylation (Fig. 5*C*).

In addition to PABP, the functional central region of DAZL (amino acids 33-190; Fig. *3A & C*) binds several other proteins such as DAZ-associated protein 1 (DAZAP1) and DAZAP2 (68), neither of which have a known role in regulating poly(A)-tail length. Pumilio also interacts with DAZL and, although the binding site has not been mapped, it is known to bind regions overlapping the RRM and DAZ motifs in DAZ and BOULE (47). However, since Pumilio has been reported to enhance deadenylation (73), we consider a role in DAZL-mediated deadenylation inhibition unlikely.

A role of DAZL in preventing deadenylation, rather than promoting polyadenylation, is entirely compatible with the data, if not the conclusions, from the zebrafish study that showed that ectopic expression of DAZL (but not DAZL mutants) in zebrafish somatic cells prevents microRNA-430 (miR430)–mediated deadenylation of an mRNA containing the *Tudor domain-containing protein 7 (tdrd7)* 3’UTR, which contains multiple miR-430 target sites as well as DAZL-binding sites (66). A mutated version of this mRNA lacking the miR-430 sites, and therefore refractory to miR-430-induced deadenylation, had a longer poly(A) tail in the presence of DAZL than in its absence, interpreted as evidence of DAZL-induced cytoplasmic polyadenylation. However, this conclusion is only valid if deadenylation of the *tdrd7* reporter mRNA is entirely miR-430 dependent, which was not demonstrated. The data can equally be explained by DAZL abrogating miR-430 independent deadenylation, which would also lead to a longer poly(A) tail on the mutated miR-430-insensitive reporter mRNA.

Consistent with the zebrafish report (66), in which apparent poly(A) extension was not dependent on efficient translation, multiple lines of evidence show that inhibition of deadenylation by the DAZL-PABP complex in our system is likewise independent of translational activation (Fig. 4*B*), and can occur even in the absence of translation (Fig. 4*C*; Fig. 5*B*). This contrasts, for instance, with PABP-dependent mRNA stabilisation in yeast which requires active translation (13). mRNA translation and stability have long been known to be intimately coupled and a theme is emerging that suggests that inefficient or repressed translation facilitates deadenylation and mRNA decay in several systems. For instance, microRNA-mediated mRNA decay requires prior translational repression that triggers CCR4-NOT-mediated deadenylation (75). Furthermore, it has recently been found that CCR-NOT4 monitors the efficiency of translation elongation through direct association with ribosomes (8), providing mechanistic insight into the accumulating evidence that inefficient translation elongation is associated with reduced mRNA stability in various organisms (4, 27, 46, 48, 51, 52). Defective translation initiation has also been shown to increase rates of deadenylation and mRNA decay (59). A recent model proposes that active translation stabilises the mRNA closed-loop structure and that the translation termination factor eukaryotic release factor (eRF)3 may compete with deadenylases for binding the PABC domain (40). Although these initiation/termination events may contribute to some instances of translation-dependent mRNA stabilisation, our data do not suggest that they operate in the protective activity of the DAZL-PABP complex because mutagenesis of M161 in PABP indicated that closed-loop formation is dispensable and, furthermore, deletion analysis showed that the PABPC domain is not required (Fig. *5A & C*). Rather, the requirement for functional PABP RRMs (Fig. 5*D*) suggests that DAZL-PABP-dependent inhibition of deadenylation may instead be dependent on competition between PABP RRMs and deadenylases for access to the poly(A) tail. An increase in the local concentration of PABP due to DAZL-mediated recruitment in addition to poly(A)-dependent binding, and/or stabilisation of the PABP-poly(A) interaction by DAZL would favour poly(A) stability by sterically occluding deadenylases (Fig. 5*E*). Although, in dissecting this mechanism, we find that translation is not required to inhibit deadenylation, it is likely that natural polyadenylated, DAZL-bound mRNAs are very actively translated (14, 64) and consequently probably stabilised by both translation-dependent and independent mechanisms. Several studies of male DAZL knockout mice note reduced stability of specific DAZL-bound mRNAs (43, 83), which is consistent with a poly(A)-protective function of DAZL since deadenylation is usually the initial step in mRNA decay (10).

Recent genome-wide studies have confirmed the coupling between poly(A)-tail length and translational efficiency in oocytes and early embryos (in contrast to somatic cells) in a variety of species including frog (18, 65). The newly defined activity of DAZL in conferring poly(A)-tail stability therefore adds to the complexity of translational control in maturing oocytes. Many verified DAZL targets of translational control (including *Dazl* mRNA) also contain CPEs and it has been proposed that initial CPEB-dependent translation of DAZL targets is reinforced by DAZL (54), synthesis of which is sustained by DAZL-mediated positive autoregulation (11). Moreover, DAZL appears to contribute to CPEB1-mediated translational regulation, perhaps by facilitating CPEB1 binding, resulting in synergism between the activity of the two proteins (64). Our results suggest that DAZL-mediated autoregulation and DAZL-CPEB1 interplay may extend further, with DAZL acting to stabilise or maintain CBEP-dependent (as well as CPE-independent) polyadenylation.

## MATERIALS AND METHODS

### Plasmids

#### Tether-function MS2-fusion plasmids

MS2-fusion plasmids pMSPN (expressing MS2 coat protein), pMS2-U1A, pMS2-PAB (PABP1), pMS2 1-4 (PABP1 RRM1-4), pMS2 1-2 (PABP1 RRM1-2), pMS2-Rd (encoding RNA binding-deficient RRMs 1 & 2), pMS2 3-4 (PABP1 RRM3-4), pMS2-Ct (PABP1 C-terminal region) (25), pMS2-mDazl, pMS2 (Δ99-166), pMS2 (33–190), pMS2-hDAZL, pMS2-hDAZ, pMS2-hBOULE (14), the construct encoding the MS2-SLBP fusion protein (23) and pMSPN-*d*0-7 (encoding MS2 fused to ICP27 amino acids 201-512 [MS2-ICP27-Ct]) (62) were previously described. pMSPN-MmPAPB1 (encoding a fusion of MS2 with murine Pabp1) was constructed by polymerase chain reaction (PCR) amplification of the full length murine *Pabp1* open reading frame (ORF) from IMAGE clone 6816124 with primers: 5’-CAG TCA GCT AGC ATG AAC CCC AGC GCC CCC-3’ and 5’-CAG TCA GGA TCC TTA GAC AGT TGG AAC ACC AGT GGC-3’. The PCR product was cut with Nhe1 and BamH1 and ligated into pMSPN which was linearised with Nhe1 and BamH1. pMSPN-MmPABP1 (M161V) was created by PCR-based site directed mutagenesis of IMAGE clone 6816124 to introduce a single point mutation into RRM2 of mouse *Pabp1* using oligonucleotides: 5’-CAC TTT ACG ATC ATT TAG AAG CAC CCC ATT CAT TTT TTC AAT AG-3’ and 5’-GAG CTA TTG AAA AAA TGA ATG GGG TGC TTC TAA ATG ATC GTA AAG TG-3’. The resulting mutation changes Pabp1 methionine 161 to valine. The entire mutant *Pabp1* (M161V) ORF was then PCR amplified with primers: 5’-CAG TCA GCT AGC ATG AAC CCC AGC GCC CCC-3’ and 5’-CAG TCA GGA TCC TTA GAC AGT TGG AAC ACC AGT GGC-3’. The PCR product was cut with Nhe1 and BamH1 and ligated into pMSPN which was linearised with Nhe1 and BamH1. pMS2 1-2-4Rd (full-length PABP1 with RNA-binding deficient RRMs 1, 2 & 4) was created by in multistep process as follows. First, RRM4 was excised from pMS2 3-4 by digestion with HindIII and AflII and replaced with a fragment created by PCR overlap-extension site-directed mutagenesis(28) using the following primers: 5’-GGT GGT CGC AGT CAA GGC TTT GGT GTT GTA TGC-3’ (4Rd1), 5’-GAC AGC TTA TCA TCG ATA AGC TTT AAT GCG-3’ (HindIIIas), 5’-GCA TAC AAC ACC AAA GCC TTG ACT GCG ACC ACC-3’ (4Rd2) and 5’-GCG AAA CTG AGC TTA AGC GCA AGT TTG AAC-3’ (AflIIs). The PCR product from primers 4Rd1 and HindIIIas and the PCR product from primers 4Rd2 and AflIIs, which contain overlapping sequence were mixed and used as template for a further PCR reaction with primers AflIIs and HindIIIas, the product of which was ligated into pMS2 3-4 cut with AflII and HindIII, resulting in pMS2 3-4Rd, encoding wildtype RRM3 and mutated RRM4 with Lys to Gln and Phe to Val at positions 1 and 5, respectively, of the octameric RNP1 motif, predicted to abolish RNA binding. Second, a BlpI fragment was excised from pMS2-PAB and ligated into pMS2-Rd linearised with BlpI, yielding pMS2-PAB-Rd1-2 (encoding fulllength PABP1 with RNA binding-deficient RRMs 1 & 2). Third, a fragment encoding mutant RRM4 was amplified from pMS2 3-4Rd with primers AflIIs and 5’-CAG TCA TGA TCA CAG GAT TTG GTA CAC GAA CAC TTG CC-3’ (4MutB), digested with AflII and BclI and ligated into pMS2-PAB-Rd1-2 digested with AflII and BclI, yielding pMS2 1-2-4Rd.

#### Tether-function reporter constructs

pLG-MS2 (expressing Luc-MS2 and Luc-MS2-cycB1), pLGENB1 (expressing Luc-ΔMS2) (25), the CSFV-IRES-Luciferase-MS2 reporter plasmid (23) and pCSFV-*lacZ* (expressing the CSFV-IRES-*lacZ* internal control mRNA) (62) were previously described. pLGMS2-ΔHex (in which the cyclin B1 3’UTR Hex sequence is mutated to AAGAAA) was created by site-directed mutagenesis of pLG-MS2 with the following oligonucleotides: 5’-GTT TTA CTG GTT TTA AGA AAG CTC ATT TTA ACA TCT AGG ATC CCC-3’ and 5’-GGG GAT CCT AGA TGT TAA AAT GAG CTT TCT TAA AAC CAG TAA AAC-3’. pLG-MS2-ΔCPE (which lacks the cyclin B1 3’UTR sequence containing the CPEs but retains the Hex sequence) was created by digesting pLG-MS2 with BamHI and BglII, removing the cyclin B1 3’UTR, and reinserting the Hex by ligating the following annealed oligonucleotides to BamHI/BglII-digested pLG-MS2: 5’-GAT CCT AGA TGT TAA TGA GCT TTA TTA ATA CAC A-3’ and 5’-GAT CTG TGT ATT AAT AAA GCT CAT TAA CAT CTA G-3’. pGEM-ΔORF-MS2 was created by PCR amplification of a fragment containing the 3 MS2-binding sites from pLG-MS2 with the following primers: 5’-TTT TGAATT CGA TCC TCG AGT CCG TTG AGA AGA AG-3’ and 5’-TTT TCA TAT GCG GGG ATC CTA GAT GTT AAA ATG AGC-3’, digesting the product with EcoRI and NdeI and ligating into ATG-pGem (a mutated version of pGEM T-easy (Promega) in which upstream ATG sequences have been removed (24)), also digested with EcoRI and NdeI. pHP-Luc-MS2 (Fig. S1 [13]) was constructed by ligating the annealed oligonucleotides 5’-AGC TCC CTG CGG TCC ACC ACG GCC GAT ATC ACG GCC GTG GTG GAC CGC AGG GAA ACA ACA ACA ACA A-3’ and 5’-CAT GTT GTT GTT GTT GTT TCC CTG CGG TCC ACC ACG GCC GTG ATA TCG GCC GTG GTG GAC CGC AGG G-3’containing a −50 kcal/mol hairpin (2) into pLG-MS2 digested with HindIII/NcoI destroying both restriction sites.

pUC19LUC-Scp3wt, encoding firefly luciferase fused to the wt Sycp3 3’UTR (Luc-Syc (Fig. S1 [10]), was previously described (54). pUC19LUC-Scp3-ΔHex, encoding Luc-Syc-ΔHex (Fig. S1 [11]) in which the Hex sequence is mutated to AAGAAA, was with the following oligonucleotides: 5’-GTT CTC TCC ACG ATT GTG TCA AGA AAG ATG ATT TAA ATT T-3’ and 5’-AAA TTT AAA TCA TCT TTC TTG ACA CAA TCG TGG AGA GAA C-3’. pUC19LUC-Scp3-Δ4-ΔHex, encoding Luc-Syc-Δ4-ΔHex (Fig. S1 [12]) in which multiple putative Dazl-binding sites and the Hex sequence are mutated, was created by sequential rounds of site-directed mutagenesis of pUC19LUC-Scp3wt with the following pairs of oligonucleotides: 1. 5’-CCA GCT ATT CCA ATG TAT CAA ACT TTC AGG GCC TCC CTC TTG TTT GTT TTA ATA GTT GTT CTC TCC-3’ and 5’-GGA GAG AAC AAC TAT TAA AAC AAA CAA GAG GGA GGC CCT GAA AGT TTG ATA CAT TGG AAT AGC TGG-3’; 2. 5’-CCA ATG TAT CAA ACT TTC AGG GCC TCC CTC CCT CCC TTT TTA ATA GTT GTT CTC TCC ACG ATT GTG TC-3’ and 5’-GAC ACA ATC GTG GAG AGA ACA ACT ATT AAA AAG GGA GGG AGG GAG GCC CTG AAA GTT TGA TAC ATT GG-3’; 3. 5’-GGG CCT CCC TCC CTC CCA TTT TAA TAG TTG TTC TCT CC-3’ and 5’-GGA GAG AAC AAC TAT TAA AAT GGG AGG GAG GGA GGC CC-3’; 4. 5’-GGC CTC CCT CCC TCC CAT TTT AAT ATT ATT TCT CTC CAC GAT TGT-3’ and 5’-ACA ATC GTG GAG AGA AAT AAT ATT AAA ATG GGA GGG AGG GAG GCC-3’; 5. 5’-TAT TAT TTC TCT CCA CGA TTG TGT CAA GAA AGA TGA TTT AAA TTT ACT AGT CAA TC-3’ and 5’-GAT TGA CTA GTA AAT TTA AAT CAT CTT TCT TGA CAC AAT CGT GGA GAG AAA TAA TA-3’.

Mutagenesis of the Tex19.1 3’UTR was carried out in pRL-TK-Tex19.1+3’UTR and pRL-TK-Tex19.1+3’UTR-Δ3 (in which one putative Dazl-binding site had been mutated) (64), kindly provided by Prof. M. Conti (University of California, San Francisco). pRL-TK-Tex19.1+3’UTR-Δ123ΔHex, in which two further putative Dazl-binding sites as well as the Hex sequence were mutated, was created by sequential mutagenesis of pRL-TK-Tex19.1+3’UTR-Δ3 with the following pairs of oligonucleotides: 1. 5’-TCA AGA TAC CTG ATT TTA GGG TTC ACT GTT TTA AAA ATT TTA CTT TGT TCG TTA CTC GTG CTC CTT TTG-3’ and 5’-CAA AAG GAG CAC GAG TAA CGA ACA AAG TAA AAT TTT TAA AAC AGT GAA CCC TAA AAT CAG GTA TCT TGA-3’; 2. 5’-CAA GAT ACC TGA TTT TAG GGT TCA CTG TTT TAA AAA TTT TAC TAA AAA CGT TAC TCG TGC TCC TTT TGA G-3’ and 5’-CTC AAA AGG AGC ACG AGT AAC GTT TTT AGT AAA ATT TTT AAA ACA GTG AAC CCT AAA ATC AGG TAT CTT G-3’; 3. 5’-GCT CCT TTT GAG GGC TTA AAA TGA CCC AAG TTG GTG TTT TGG ATC CAG GT-3’ and 5’-ACC TGG ATC CAA AAC ACC AAC TTG GGT CAT TTT AAG CCC TCA AAA GGA GC-3’. pRL-TK-Tex19.1+3’UTR-ΔHex, in which only the Hex sequence was mutated, was created by mutagenesis of pRL-TK-Tex19.1+3’UTR with the following pair of oligonucleotides: 5’-CTC CTT TTG AGG GCT TTT GTT GAC CCA AGT TGG TGT TTT GGA TCC AGG T-3’ and 5’-ACC TGG ATC CAA AAC ACC AAC TTG GGT CAA CAA AAG CCC TCA AAA GGA G-3’. Wildtype and mutated Tex19.1 3’UTRs were PCR amplified with the primers 5’-GCT AGT CGA CTG CAC ATT CCT GAG ACA C3’ and 5’-GCT AAA GCT TCT GGA TCC AAA ACA CCA AC-3’, digested with SalI/HindIII and cloned downstream of the firefly luciferase ORF in pUC19LUC (55), also digested with SalI/HindIII generating pUC19LUC-Tex19wt, pUC19LUC-Tex-ΔHex and pUC19LUC-Δ123-ΔHex, used to generate reporter mRNAs (Fig. S1 [7], [8] & [9]).

### Site-directed mutagenesis

Site-directed mutagenesis was carried out with a QuickChange^®^ Lightning Site-Directed Mutagenesis Kit (Agilent) according to manufacturer’s instructions. All mutations were verified by DNA sequencing.

### *In vitro* transcription, polyadenylation and cordycepin incorporation

Plasmids were linearised with appropriate restriction enzymes and T7- or Sp6- (for internal control mRNA) dependent *in vitro* transcription was performed as previously described (25). Luciferase reporter mRNAs were m^7^GpppG-capped except those containing an IRES or when specifically stated (ApppG). *In vitro* polyadenylation was carried out with a Poly(A)-tailing Kit (Ambion) or with *E. coli* poly(A) polymerase (New England Biolabs) according to manufacturer’s instructions. Cordycepin was added to the 3’ end of *in vitro* transcribed mRNA by substituting 50 mM Cordycepin (3’-deoxyadenosine) 5’-triphosphate sodium salt (Sigma-Aldrich) in 10 mM Tris pH7 for ATP in a poly(A)-tailing reaction with a Poly(A) Tailing Kit (Ambion) as described above. All *in vitro*-generated mRNAs were purified by acid phenol/chloroform (Ambion) extraction, removal of unincorporated nucleotides with a Chromaspin column (Clontech) and ethanol precipitation.

### Oocyte isolation, tether-function analysis and metabolic labelling

*X. laevis* oocytes (stage VI) were isolated from excised ovary by manual dissection or collagenase treatment and maturation was induced with 10 μg/ml progesterone as previously described (63). Tethered-function assays and verification of expression of MS2-fusion proteins by metabolic labelling were performed as described previously (63). Oocytes were injected with 50 nl mRNA encoding MS2-fusion proteins at 1 μg/μl and reporter/control mRNA at 10 ng/μl, except for RNA coimmunoprecipitation (0.5 ng/μl). Analysis of luciferase and β-galactosidase activity was previously described (25).

### RNA extraction from oocytes

Pooled oocytes (20–30) were homogenised in 1 ml Tri Reagent (Sigma-Aldrich), centrifuged at 20000 g for 10 min, and the supernatant was mixed with 200 μl chloroform. Phases were separated by centrifugation (20000 g, 10 min) and RNA was precipitated from the aqueous phase with 500 μl isopropanol, followed by centrifugation (20000 g, 15 min). The pellet was washed with 80% ethanol, briefly dried and resuspended in 100 μl RNase-free water. The RNA solution was further purified with an RNeasy Mini Kit (Qiagen) following manufacturer’s instructions. The RNA was eluted in 30 μl RNase-free water, the concentration determined with a NanoDrop™ One spectrophotometer (ThermoFisher Scientific), and stored at −80°C until further use.

### Poly(A) analysis

RNA ligation-coupled PCR was modified from (9). 6 μg total oocyte RNA, from pools of 20 to 30 oocytes, was ligated to 60 pmol P1 anchor primer (5’-phosphate-GGT CAC CTT GAT CTG AAG C-3’-amino modifier), in a 10 μl reaction using 10 U T4 RNA ligase (New England Biolabs) in 50 mM Tris-HCl (pH 7.5), 10 mM MgCl_2_, 1 mM DTT, 1 mM ATP for 1 hr at 37°C, followed by thermal enzyme inactivation (2 min, 95°C). 50 pmol reverse transcription primer (P1’, complementary to P1; 5’-GCT TCA GAT CAA GGT GAC C-3’) was added to the ligation mix and hybridised by heating to 65°C for 5 min and rapidly cooling on ice. Reverse transcription was performed in a 50 μl reaction using 200 U Superscript III (ThermoFisher Scientific) in FS buffer (50 mM Tris-HCl (pH 8.3), 75 mM KCl, 3mM MgCl_2_) supplemented with 5 mM DTT, 10 U RNasin ribonuclease inhibitor (Promega) and 0.5 mM each of dATP, dCTP, dGTP and dTTP (dNTPs) at 55°C for 1 hr, followed by thermal enzyme inactivation (15 min, 70°C). 1 μl of the cDNA preparation was amplified in a 50 μl PCR reaction using 1.25 U Dream Taq (ThermoFisher Scientific) DNA polymerase in its supplied buffer with 0.2 mM dNTPs and 0.2 μM P1’ and gene-specific primers (MS2: 5’-CTC TCT CTC AGG GCT GAT TAC TAG-3’; *Sycp3:* 5’-ACC AGC TAT TCC AAT GTA TCA AAC-3’; *Tex19.1:* CTG TCT TAG GGT CAA GAT ACC-3’) for 35-40 cycles (95°C, 30 s; 56°C, 1 min; 72°C, 1 min). Products were analysed by electrophoresis on a 2% agarose gel.

### RNase H treatment

6 μg total oocyte RNA from pooled injected oocytes was treated with 10 U RNase H (Ambion) and 500 ng oligo(dT) in 20 mM HEPES-KOH (pH 8.0), 50 mM KCl, 4 mM MgCl_2_, 1mM DTT, 50 μg/ml BSA at 37°C for 30 min. The reaction was stopped by extraction with acidic phenol/chloroform and the RNA was ethanol precipitated in the presence of 0.3 M sodium acetate (pH 5.2) and resuspended in RNase-free water.

### RNA Immunoprecipitation

For RNA co-immunoprecipitation, oocytes were microinjected with 50 ng MS2 or MS2-DAZL mRNA and, after 6 hr incubation, again with 25 pg ApG-capped, non-adenylated luciferase reporter mRNA (Fig. S1 [4]) and incubated for 3 hr. Oocytes were homogenised in 10 μl/oocyte of 20 mM Hepes pH 7.6, 100 mM NaCl, 1.5 mM MgCl_2_, 10 mM KCl, 1 mM DTT, 5mM NaF, 0.5 U/μl RNasin (Promega) 0.5% Triton X-100, 0.5% Na deoxycholate in the presence of protease inhibitor cocktail (Roche). ePABP was immunoprecipitated with 0.5 μg anti-ePABP antibody/mg total protein (or rabbit IgG as a control) in the presence of 50 μl protein A-sepharose for 2 hr at 4°C. Beads were washed 7 times in 20 mM Hepes pH 7.6, 10 mM KCl, 1 mM MgCl_2_, 350 mM NaCl, 0.05% NP-40. Samples were evenly split for protein and RNA analysis. Protein was analysed by immunoblotting. RNA was purified using Tri Reagent (Sigma) and analysed by 25-cycle amplification using Titan One Tube RT-PCR (Roche) with primers 5’-GGC GCG GTC GGT AAA GTT-3’ and 5’-AGC GTT TTC CCG GTA TCC A-3’ (luciferase) or 5’-TGG TGA CAA TGC CAT GTT C-3’ and 5’-GAT AAT GGA TCT GGT ATG TG-3’ (β-actin).

### Immunoblotting

Oocyte lysates or immunoprecipitates were resolved on 4-12% NuPAGE gels (Invitrogen) and transferred to polyvinylidene fluoride membranes (Millipore). Membranes were probed with anti-gePABP (1:2000) primary antibodies (77) and horseradish peroxidase-conjugated goat anti-rabbit IgG secondary antibodies (Sigma-Aldrich) (1:50000), and detected by enhanced chemiluminescence (ThermoFisher).

## ACKNOWLEDGEMENTS

We thank Prof. Marco Conti (University of California, San Francisco) for kind provision of plasmids and Matthew Brook for critical reading of the manuscript. This work was funded by the Wellcome Trust and the Medical Research Council. This work was undertaken in the Medical Research Council Centre for Reproductive Health, which is funded by Medical Research Council Centre grant MR/N022556/1.

## LEGENDS TO SUPPORTING FIGURES

**Fig. S1.**
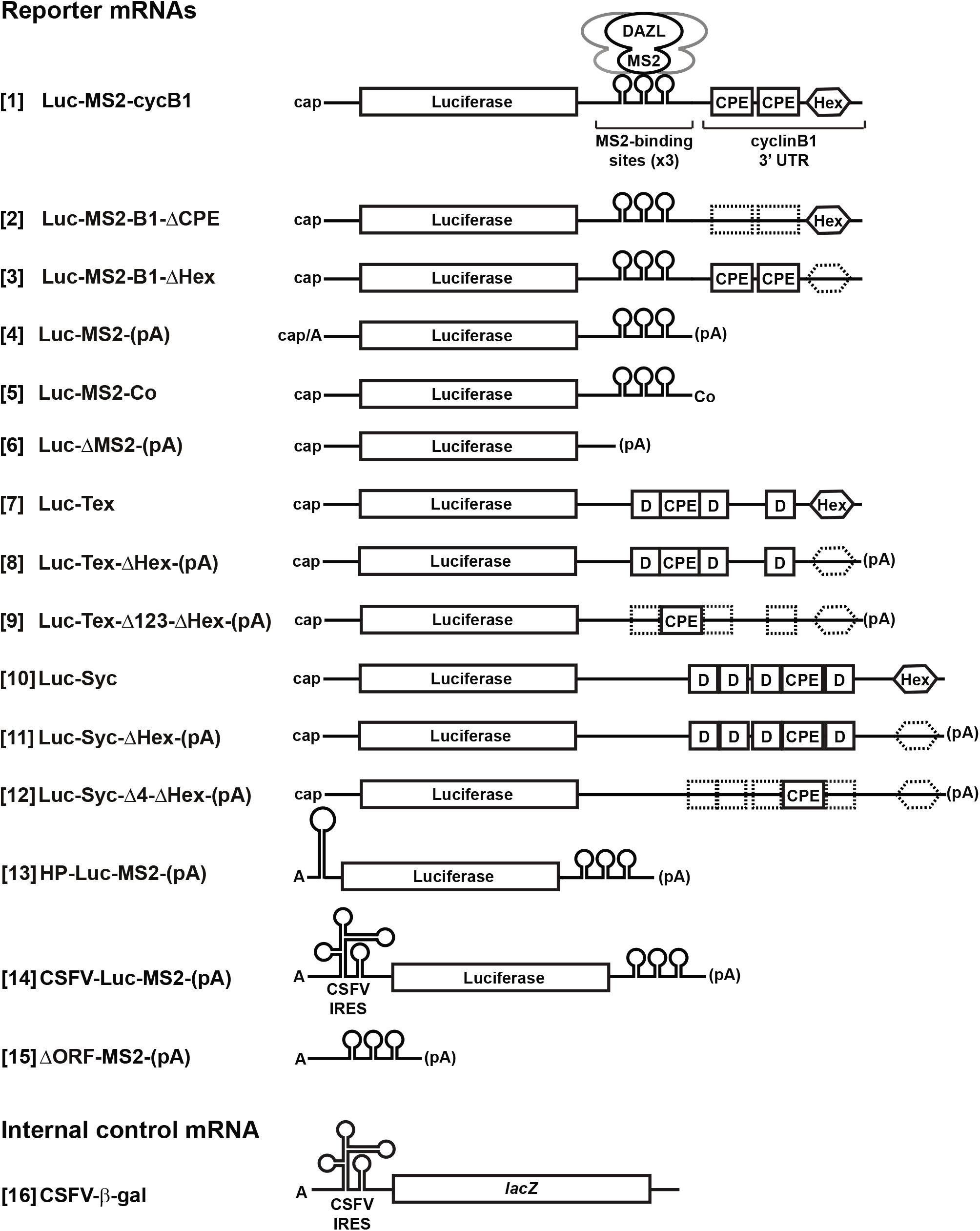
Diagram of reporter and internal control mRNAs used in the tether-function assay and analysis of natural DAZL target mRNAs. DAZL is expressed in *X. laevis* stage VI oocytes as a fusion with the phage MS2 coat protein (MS2) and tethered to various luciferase reporter mRNAs via the interaction of MS2 with three cognate binding sites within the 3’UTR. Reporter mRNAs are unadenylated or polyadenylated (pA) as indicated, and have a 5’ m^7^GpppG or ApppG cap (indicated by ‘cap’ or ‘A’, respectively). Reporter mRNAs carrying a wildtype [1] or mutant [2, 3] cyclin B1 3’ UTR (‘cycB1’) were unadenylated when injected. Cap- and eIF4F-independent initiation of translation is mediated by the classical swine fever virus (CSFV) internal ribosome entry site (IRES) and mRNAs carrying this sequence element [14, 16] were ApppG-capped. A luciferase reporter mRNA carrying a 3’ deoxyadenosine (cordycepin) is indicated by ‘Co’ [5] and an mRNA lacking the MS2-binding sites is designated Luc-ΔMS2 [6]. For luciferase assays, an ApppG-capped, unadenylated, CSFV IRES-dependent internal control mRNA expressing β-galactosidase was used [16]. D, putative DAZL-binding site (UUGUU); CPE, cytoplasmic polyadenylation element; Hex, polyadenylation signal hexanucleotide. Mutated sequence elements are indicated by dashed boxes.

**Fig. S2.**
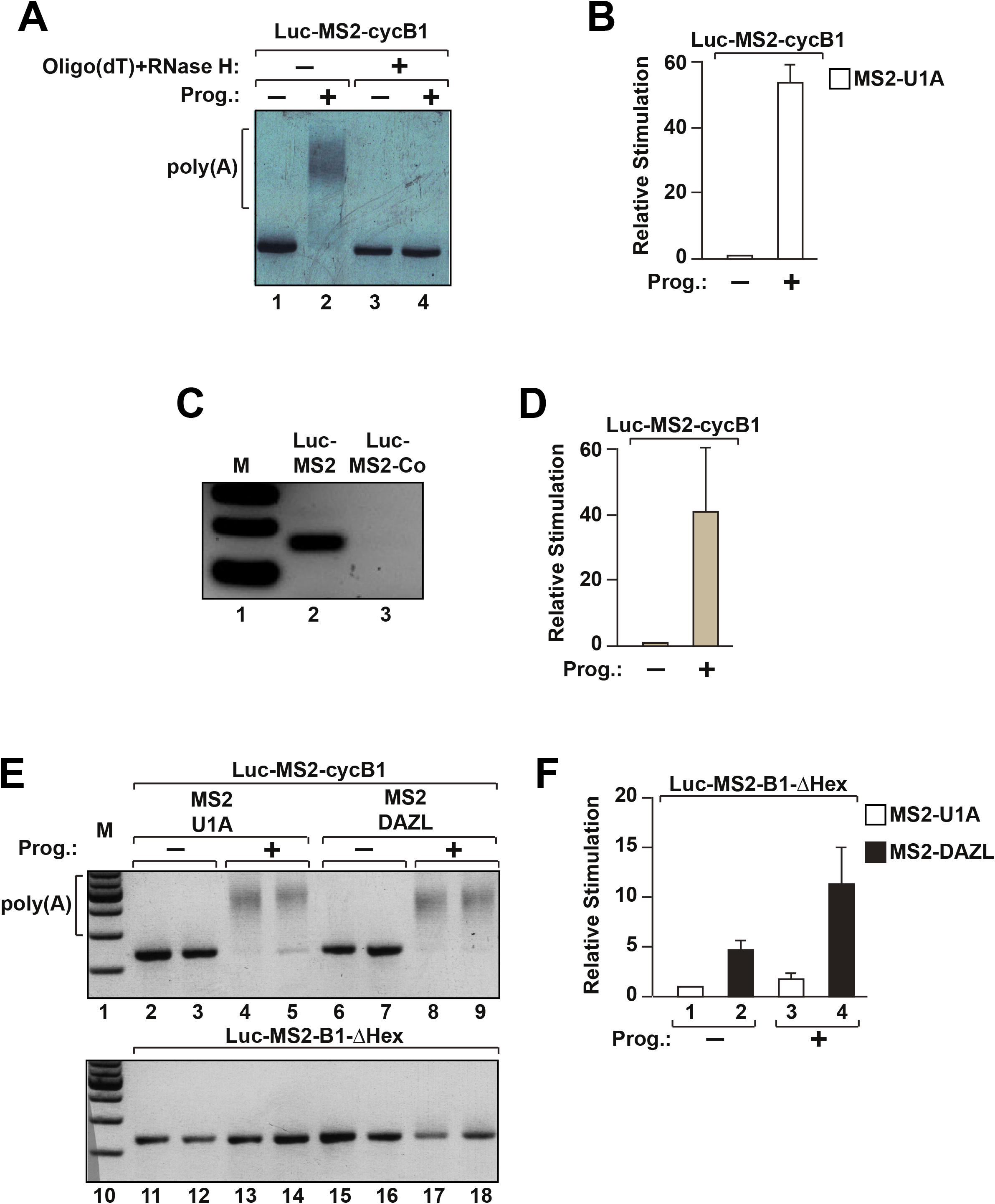
Polyadenylation is not required for DAZL-mediated translational activation, nor is it induced by DAZL. ***(A)*** Oocytes were injected with Luc-MS2-cycB1 (Fig. S1 [1]) and treated with progesterone (Prog.) (+) or left untreated (-). mRNA was extracted and either treated with oligo(dT) and RNase H (+) or left untreated (-), followed by a PAT assay and gel electrophoresis. This demonstrates that the high molecular weight bands (indicated by ‘poly(A)’) are due to polyadenylation. **(*B*)** Effects of progesterone-induced maturation on translation of Luc-MS2-cycB1 were measured by luciferase assay normalised to β-galactosidase activity of a coinjected CSFV-β-gal internal control mRNA (Fig. S1 [16]). Injected oocytes were treated with progesterone (Prog.) (+) or left untreated (-). Translational stimulation relative to Luc-MS2-cycB1 in untreated cells (set to 1) is plotted, showing that progesterone-induced polyadenylation strongly stimulates Luc-MS1-cycB1 translation, as expected. Error bars represent standard error of the mean (SEM); *n=3.* **(*C*)** Control experiment to verify incorporation of 3’ deoxyadenosine (cordycepin) in the reporter mRNA. Successful incorporation should prevent RNA ligation of an oligonucleotide to the 3’ end of the mRNA, the first step in a PAT assay. Equal amounts of Luc-MS2 and Luc-MS2-Co (Fig. S1 [4] and [5]) were subjected to PAT assay with a 20-cycle PCR reaction. The absence of a product with Luc-MS2-Co is indicative of cordycepin incorporation. ***(D)*** As a control for successful oocyte maturation in Fig. 1*C*, oocytes were injected in parallel with Luc-MS2-cycB1 and CSFV-β-gal mRNAs and treated with progesterone (Prog.) (+) or left untreated (-). Effects on translation were measured by luciferase assay and values were normalised to β-galactosidase activity. Translational stimulation relative to Luc-MS2-cycB1 in untreated cells (set to 1) is plotted, showing that progesterone treatment strongly stimulates Luc-MS1-cycB1 translation, indicative of successful maturation. Error bars represent SEM; *n*=3. ***(E)*** Oocytes expressing MS2-U1A or MS2-DAZL were co-injected with Luc-MS2-cycB1 or Luc-MS2-B1-ΔHex and CSFV-β-gal mRNAs (Fig. S1 [1], [3] & [16]) and treated with progesterone (Prog.) (+) or left untreated (-). Poly(A) status was analysed by PAT assay and gel electrophoresis. M, DNA size markers; poly(A), high molecular weight bands indicative of polyadenylation. Duplicate samples are shown in adjacent lanes as indicated. This shows that deletion of the Hex sequence prevents mRNA polyadenylation, as expected. ***(F)*** Effects of MS2-U1A and MS2-DAZL on translation of Luc-MS2-B1-ΔHex (progesterone-treated or untreated; see *(E))* were measured by luciferase assay normalised to β-galactosidase activity. Translational stimulation relative to Luc-MS2-cycB1 in untreated MS2-U1A-expressing cells (column 1; value set to 1) is plotted, showing that DAZL stimulates mRNA translation in the absence of polyadenylation. Error bars represent SEM; *n=3*.

**Fig. S3.**
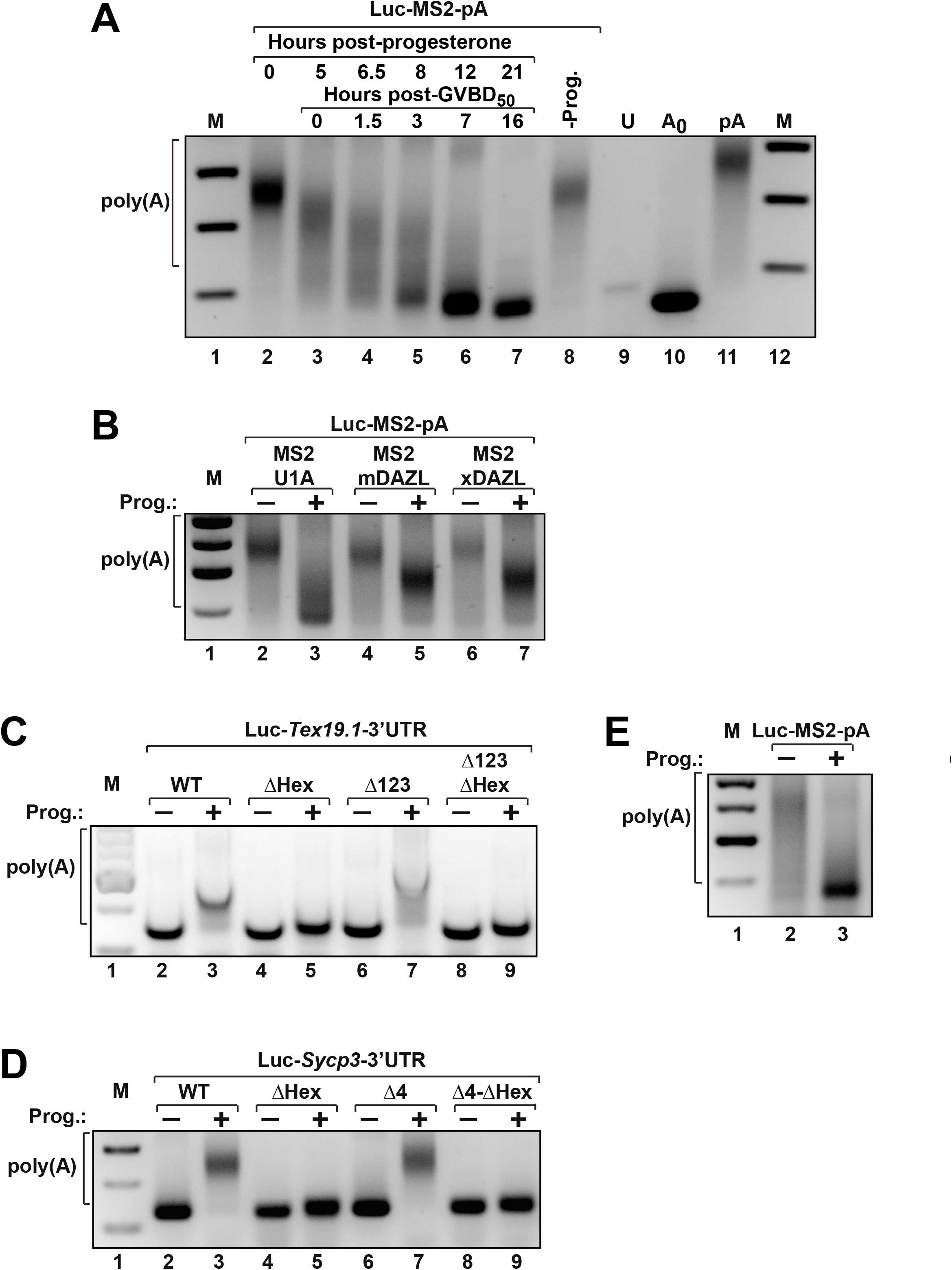
DAZL inhibits mRNA deadenylation when tethered and on natural target mRNAs. **(*A*)** Oocytes were injected with an *in vitro* polyadenylated reporter mRNA (Luc-MS2-pA; Fig. S1 [4]), treated with progesterone, harvested at the indicated times post-treatment and reporter polyadenylation analysed by PAT assay. - Prog., untreated; U, uninjected control. GVBD50 (visible breakdown of 50% of germinal vesicles) occurred at 5 hours. M, DNA size markers; poly(A), high molecular weight bands indicative of polyadenylation. **(*B*)** Oocytes expressing MS2-U1A, or MS2-murine (m)DAZL or MS2-frog (x)DAZL were co-injected with Luc-MS2-pA mRNA (Fig. S1 [4]) and treated with progesterone (Prog.) (+) or left untreated (-). mRNA polyadenylation was analysed by PAT assay and gel electrophoresis. **(*C*** & ***(D)*** Control experiments showing that mutation of the polyadenylation sequence (Hex) abolishes maturation-induced polyadenylation of reporter mRNAs containing the *Sycp3* (*C*) or *Tex19.1 (D)* 3’UTRs. Oocytes were injected with the indicated reporter mRNAs (see Fig. S1 [7]-[12]) and treated with progesterone (Prog.) (+) or left untreated (-). mRNA polyadenylation was analysed by PAT assay and gel electrophoresis. ***(E)*** Luc-MS2-pA was co-injected with Luc-Syc-ΔHex-pA (see Fig. 2*C*), treated with progesterone (Prog.) (+) or left untreated (-). Deadenylation of Luc-MS2-pA was analysed by PAT assay and gel electrophoresis, showing that this was more efficient than that of Luc-Syc-ΔHex-pA.

**Fig. S4.**
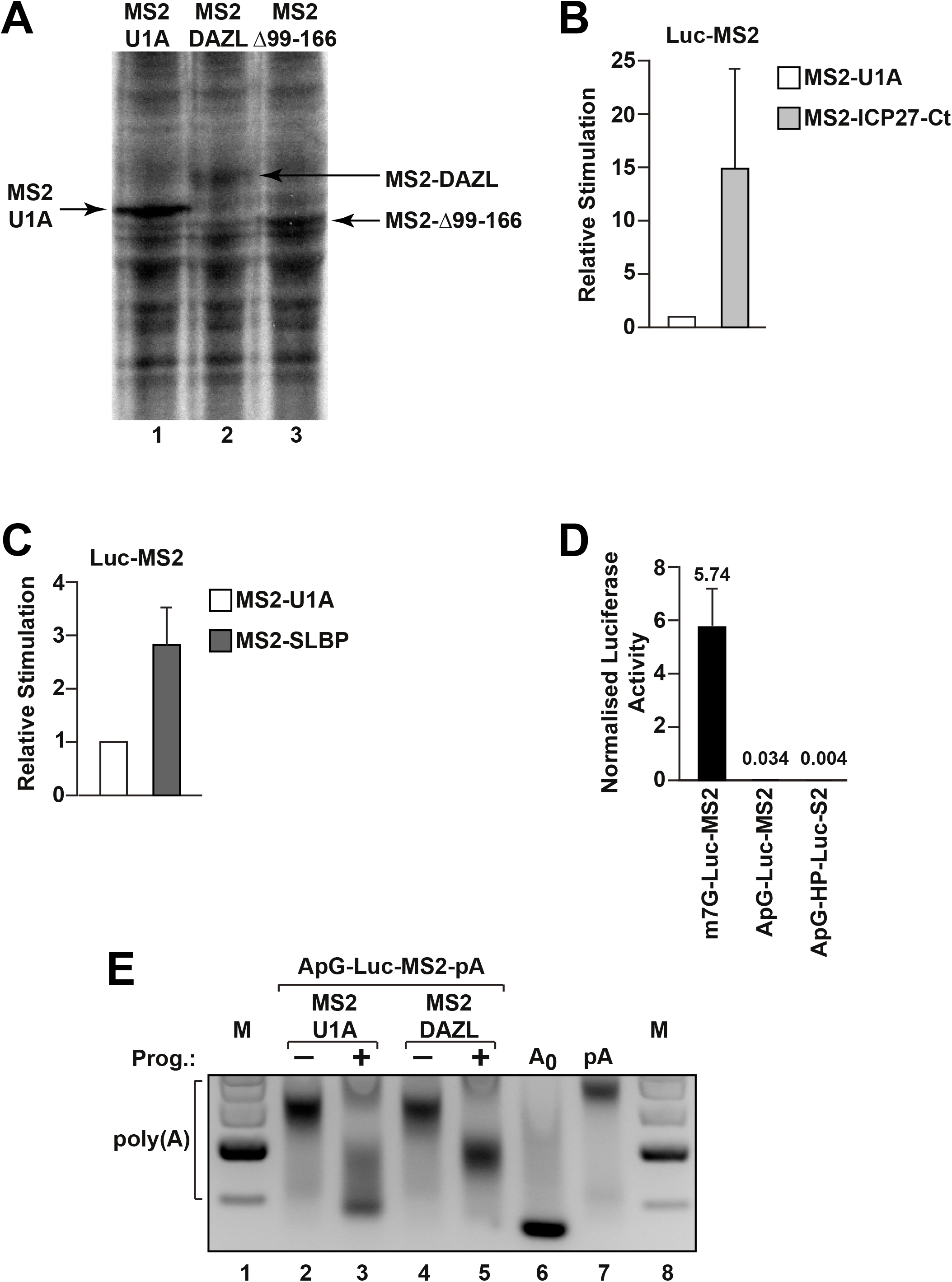
DAZL-mediated inhibition of deadenylation requires the PABP-binding site and is independent of translation. ***(A)*** Control experiment showing that MS2-DAZL (Δ99-166) is expressed in oocytes. Oocytes were injected with MS2-U1A, MS2-DAZL or MS2-DAZL (Δ99-166) mRNAs and labelled with ^35^S-Met. Total protein was extracted and analysed SDS-PAGE and autoradiography. The position of fusion proteins is indicated by arrows. MS2-U1A, MS2-DAZL and MS2-DAZL (Δ99-166) proteins contain 21, 10 and 8 methionine residues, respectively, accounting for the greater intensity of the MS2-U1A band. ***(B)*** Oocytes expressing MS2-U1A or MS2-ICP27-Ct (the C-terminal PABP-binding region of the HSV-1 ICP27 protein) were co-injected with Luc-MS2 and CSFV-β-gal mRNAs and treated with progesterone. Effects on translation were measured by luciferase assay and values normalised to β-galactosidase activity. Translational stimulation relative to that by MS2-U1A (set to 1) is plotted, showing that ICP27-Ct strongly stimulates translation in mature oocytes. Error bars represent SEM; *n*=2. **(*C*)** Oocytes expressing MS2-U1A or MS2-SLBP (stem-loop binding protein) were co-injected with Luc-MS2 and CSFV-β-gal mRNAs (Fig. S1 [4] & [16]) and treated with progesterone. Effects on translation were measured by luciferase assay and values normalised to β-galactosidase activity. Translational stimulation relative to that by MS2-U1A (set to 1) is plotted, confirming that SLBP stimulates translation in mature oocytes (1). Error bars represent SEM; *n*=3. **(*D*)** Oocytes were co-injected with the indicated reporter mRNAs (Fig. S1 [4] & [13]) and CSFV-β-gal mRNA. Effects on translation were measured by luciferase assay and values normalised to β-galactosidase activity. Values are shown above the bars for clarity; error bars represent SEM; *n*=2. ***(E)*** Oocytes expressing MS2-U1A or MS2-DAZL were co-injected with ApG-Luc-MS2-pA and treated with progesterone (Prog.) (+) or left untreated (-). mRNA polyadenylation was analysed by PAT assay and gel electrophoresis. A_0_ & pA, polyadenylation controls: PAT assays of unadenylated (A0) and *in vitro* polyadenylated (pA) reporter mRNA.

**Fig. S5.**
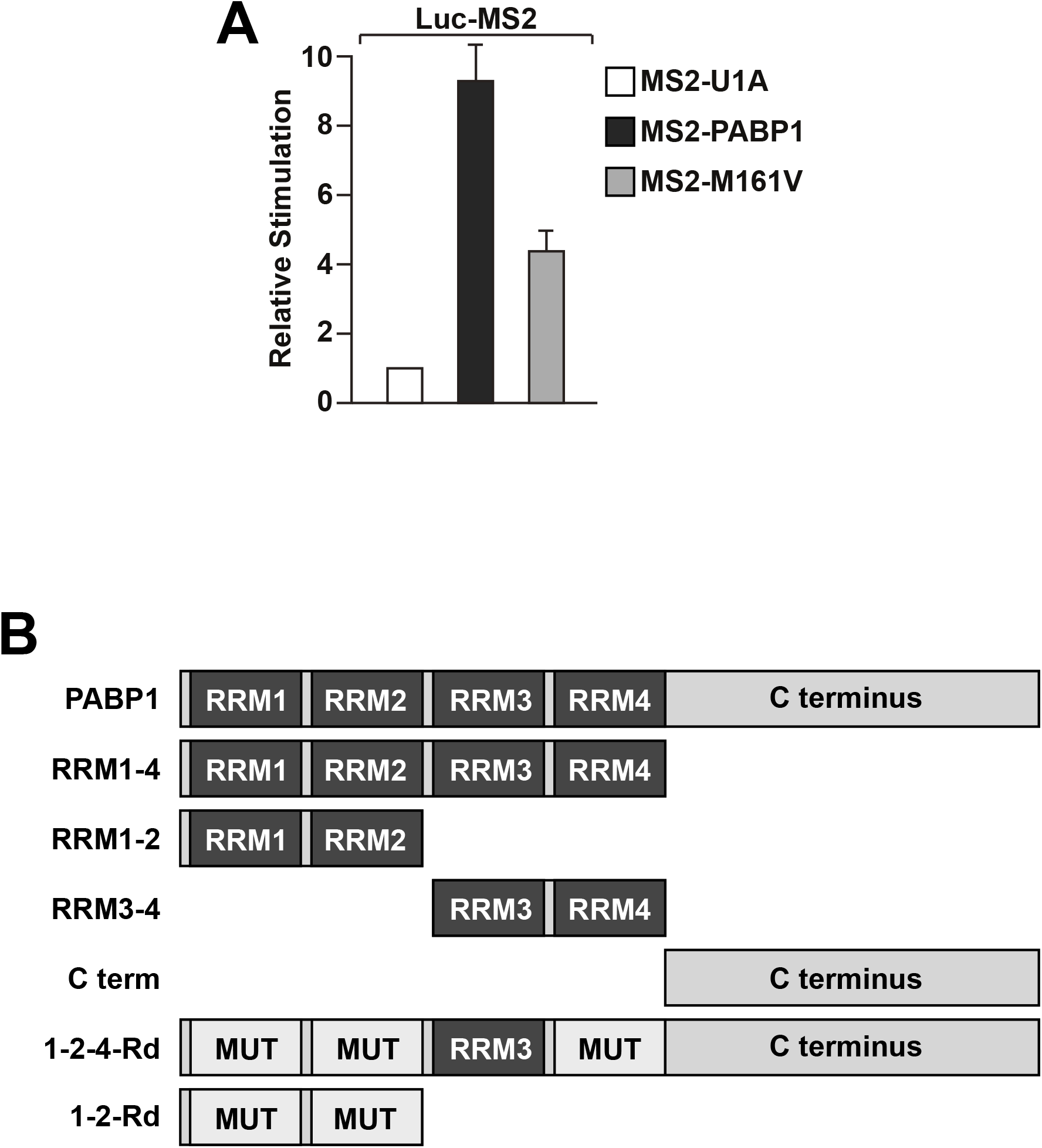
Regulation of deadenylation does not require mRNA closed-loop formation but does require functional PABP RRMs. **(*A*)** Oocytes expressing MS2-U1A, MS2-PABP1 or MS2-PABP1 (M161V) were co-injected with Luc-MS2 and CSFV-β-gal mRNAs. Effects on translation were measured by luciferase assay and values normalised to β-galactosidase activity. Translational stimulation relative to by MS2-U1A (set to 1) is plotted, confirming that the M161V mutation strongly reduces the ability of tethered PABP1 to activate translation. Error bars represent SEM; *n*=4. **(*B*)** PABP1 comprises four non-identical RNA-recognition motifs (RRMs) with different RNA-binding specificities and a C-terminal region (C terminus) that does not bind RNA (2). Full-length PABP1 and the indicated domains in isolation were used in tether-function assays described in Fig. *5B* & *C*. ‘Mut’ indicates mutated RRMs, in which the RNA-binding activity has been specifically inactivated (2).

---

**Table S1.**
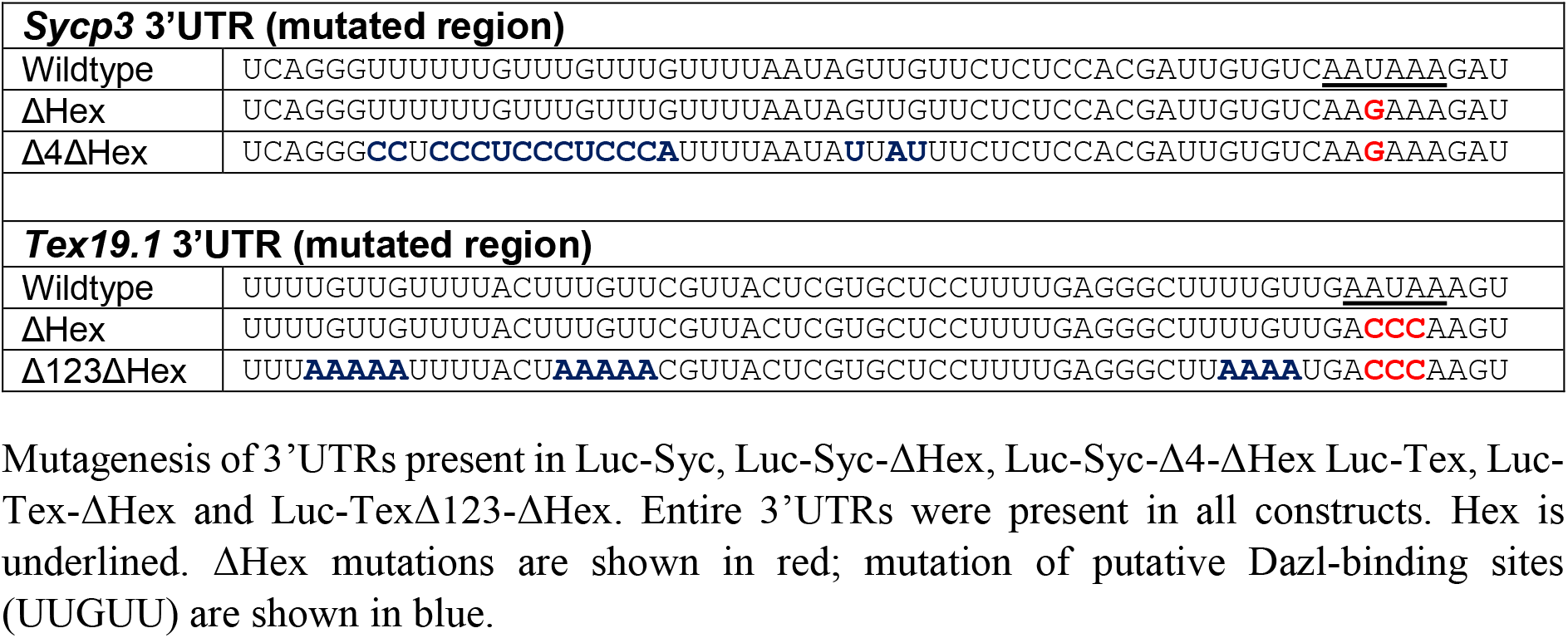
Mutagenesis of *Sycp3* and *Tex19.1* 3’UTRs.

